# Zinc pyrithione is a potent inhibitor of PL^Pro^ and cathepsin L enzymes with *ex vivo* inhibition of SARS-CoV-2 entry and replication

**DOI:** 10.1101/2022.03.03.482819

**Authors:** Jerneja Kladnik, Ana Dolinar, Jakob Kljun, David Perea, Judith Grau-Expósito, Meritxell Genescà, Marko Novinec, Maria J. Buzon, Iztok Turel

## Abstract

As SARS-CoV-2 triggered a global health crisis, there is an urgent need to provide patients with safe, effective, accessible, and preferably oral therapeutics for COVID-19 that complement mRNA vaccines. Zinc compounds are widely known for their antiviral properties. Therefore, we have prepared a library of zinc complexes with pyrithione (1-hydroxy-2(1*H*)-pyridinethione) and its analogues, all of which showed promising *in vitro* inhibition of cathepsin L, an enzyme involved in SARS-CoV-2 entry, and PL^Pro^, an enzyme involved in SARS-CoV-2 replication both in (sub)micromolar range. Zinc pyrithione **1a** is a well-established, commercially available antimicrobial agent and was therefore selected for further evaluation of its SARS-CoV-2 entry and replication inhibition in an *ex vivo* system derived from primary human lung tissue. Our results suggest that zinc pyrithione complex **1a** provides a multitarget approach to combat SARS-CoV-2 and should be considered for repurposing as a potential therapeutic against the insidious COVID-19 disease.

**Featured image:** In our study, we show that zinc pyrithione holds immense potential for the development of a possible out-patient treatment for SARS-CoV-2 due to its inhibition of viral entry and replication.

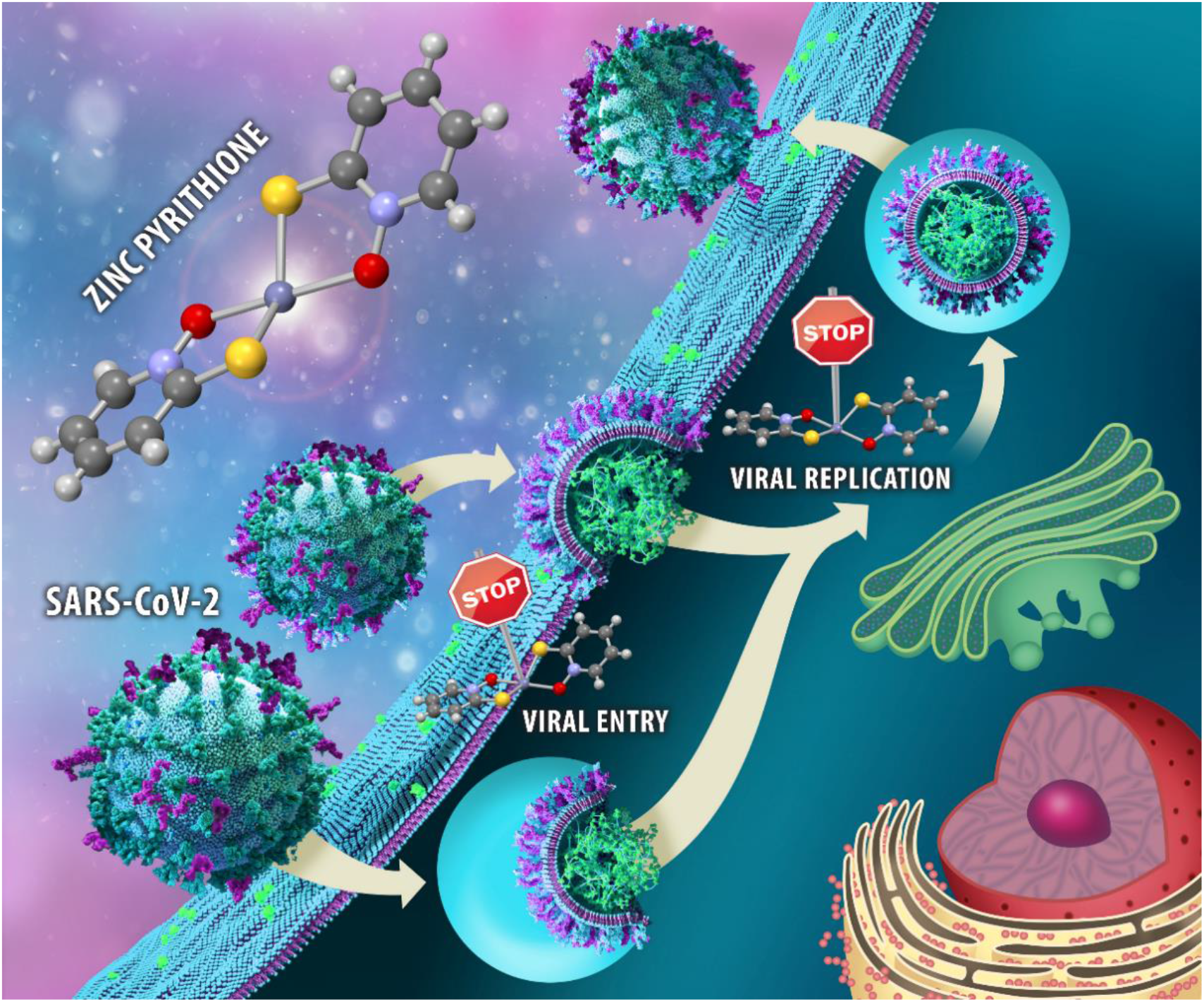

## Introduction

Research and development of safe and effective pharmacotherapy for a life-threatening COVID-19 disease must now focus on curative approaches after the rapid and successful development of mRNA vaccines. The development of specific SARS-CoV-2 inhibitors requires a thorough knowledge of viral structure, host cell entry, and the replication cycle.^1^ SARS-CoV-2 is an enveloped single-stranded positive-sense RNA virus^2^ that requires binding of the viral spike protein to the human zinc metalloproteinase receptor ACE2 as the first step in internalization process.^3^ After binding, viral entry is possible through i) direct fusion with the cell surface involving the proteases furin and transmembrane serine protease-2 (TMPRSS2), followed by direct release of SARS-CoV-2 RNA and/or ii) receptor-mediated endocytosis, in which the lysosomal cysteine protease cathepsin L facilitates the release of the genetic material after fusion of the viral and endosomal membranes.^4^ Once RNA is released into the cytosol, translation and replication of the genome occur, including the potential targets papain-like protease (PL^Pro^), chymotrypsin-like protease (3CL^Pro^, also abbreviated as M^Pro^), and RNA-dependent RNA polymerase (RdRp). After the synthesis of the viral structural proteins and their transition through endoplasmatic-reticulum-to-Golgi intermediate compartment, the viral copies are released from host cells by exocytosis.^5^

To our knowledge, the following intravenous/subcutaneous treatments are currently approved by EMA and FDA: remdesivir targeting RdRp, and antibodies targeting the spike protein, i.e., casirivimab/imdevimab, regdanvimab, sotrovimab and bamlanivimab/etesevimab, as well as tocilizumab as an interleukin-6 blocker. In addition, anakinra, a subcutaneously applied interleukin-1 receptor antagonist and orally administered PF-07321332 (SARS-CoV-2-M^Pro^ inhibitor) along with ritonavir (which slows down the metabolism of PF-07321332) were also authorized for COVID-19 treatment. Currently, molnupiravir (a ribonucleoside analogue that inhibits SARS-CoV-2 replication) and baricitinib (an immunosuppressant) are awaiting the green light for their marketing authorization.^6,7^ However, due to constant viral mutations, it is necessary to continue research and development of new therapeutic options with other targets. To skip the rigorous and lengthy process from potential drug candidate to its clinical approval, many investigations have been conducted to repurpose already approved drugs for COVID-19 indication.^8^

Zinc is the second most abundant d-block metal in the human body and plays several important roles in biological systems (e.g., structural, catalytic, regulatory, signalling).^9^ Zinc received much attention during the pandemic because of its antiviral, anti-inflammatory, and antioxidant properties.^10^ In addition, hypozincemia has also been suspected to cause dysgeusia and anosmia, common COVID-19 symptoms, because taste and smell functions are regulated by the zinc-containing enzyme carbonic anhydrase.^11,12^ Many studies suggest that zinc deficiency increases susceptibility to infectious diseases, including COVID-19.^13–16^ There is some evidence that Zn^2+^ can indirectly inhibit ACE2 receptors^10,17,18^ and furin was also reported to be inhibited by zinc.^19^ Low bioavailability of zinc may also negatively affect the activity of signalling molecules (interferon type 1, tumor necrosis factor, interleukin IL-6) leading to cytokine storm and severe lung damage.^18^

A computational study has confirmed possible inhibition of RdRp and M^Pro^ by zinc ions.^20,21^ Additionally, the crystal structure of SARS-CoV-2 M^Pro^ has confirmed the interactions of Zn^2+^ in the active site.^22,23^ Although the structural studies clearly confirmed that a zinc ion inhibits SARS-CoV-2 M^Pro^ by binding to the active site, it was also found that the experimentally determined affinity between the protein and the metal ion does not appear to be high enough for any real therapeutic applications of zinc supplementation, at least in its ionic form.^22^

In 2010, te Velthius *et al*. showed that the combination of Zn^2+^ ions and pyrithione **a** (1-hydroxy-2(1*H*)-pyridinethione or 2-pyridinethiol *N*-oxide, Fig. 1) successfully inhibited SARS-CoV replication, possibly by Zn^2+^ suppression of RdRp activity. Inhibition of SARS-CoV was less potent with pyrithione itself and even less effective with zinc acetate alone.^24^ Zinc pyrithione **1a** has also been shown to effectively inhibit M^Pro^ and PL^Pro^ of SARS-CoV^25–27^ as well as M^Pro^ of SARS-CoV-2.^28–30^ There are several reports that zinc pyrithione damages iron–sulfur (Fe–S) clusters and consequently elicits antimicrobial activity.^31–33^ Recently, Maio *et al*. suggested that Fe–S clusters act as cofactors of the SARS-CoV-2 RdRp enzyme and therefore represent another potential target to combat COVID-19.^34^

**Fig. 1:**
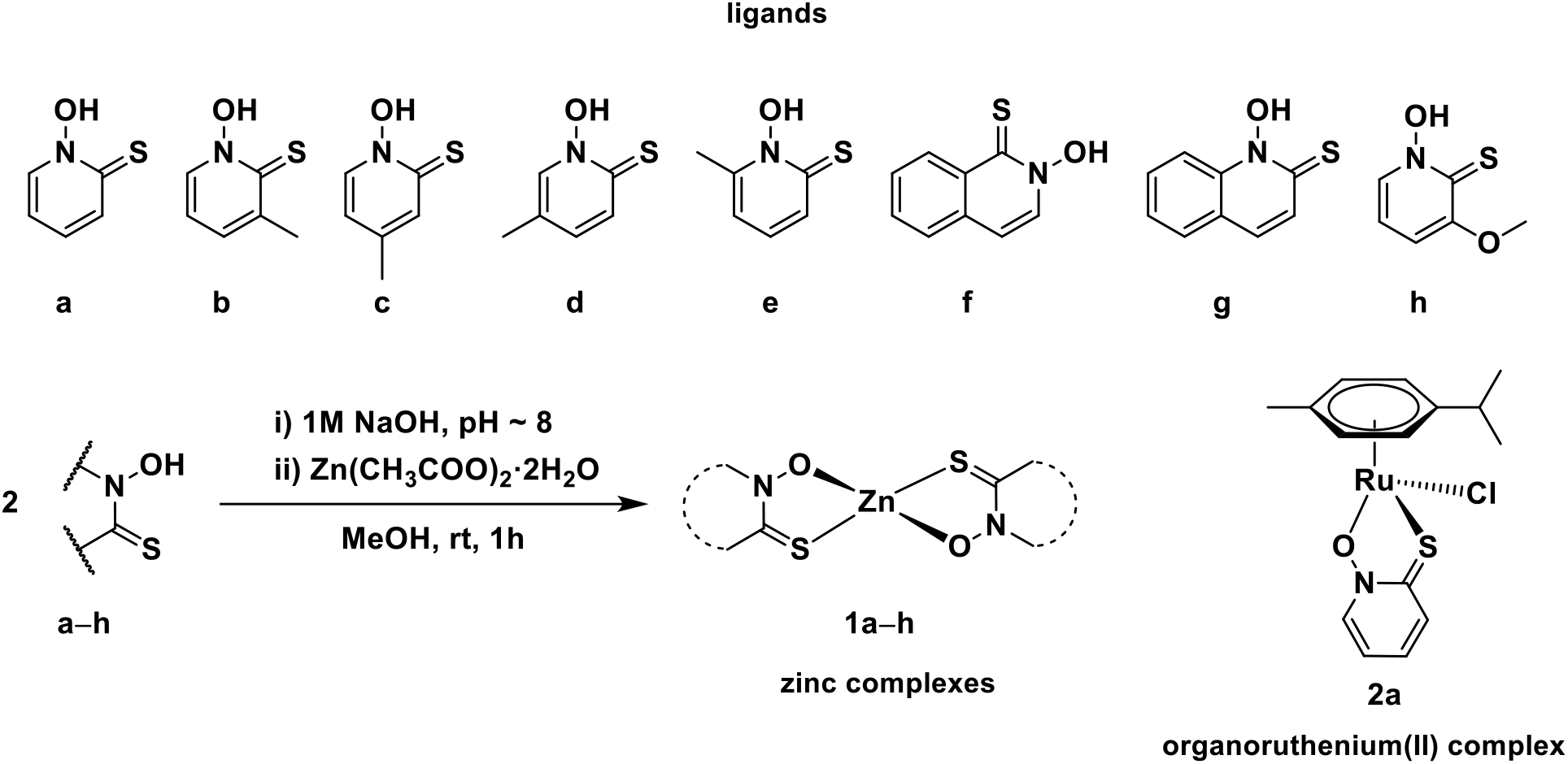
Chemical structures of investigated pyrithionato ligands a–h, synthesis of zinc complexes 1a–h and organoruthenium complex 2a. Zinc complexes **1a–h** were prepared by a modified published procedure.^41^ First, the corresponding pyrithione ligand **a–h** was dissolved in methanol. Before the addition of zinc acetate, the pyrithione ligand had to be deprotonated, which was achieved with an aqueous solution of 1M NaOH. After addition of zinc acetate to this solution, a white precipitate was obtained immediately, which was then filtered off and dried overnight in an oven. All compounds prepared were physico-chemically characterized by NMR (Supplementary Figs. 2–9), high resolution mass spectrometry, elemental analysis (C, H, N), IR spectroscopy and new crystal structures were also obtained for compounds **1d** and **1f** (see below).

Based on the described data from the literature, we decided to further investigate a system of zinc and pyrithione as a promising antiviral combination against SARS-CoV-2. Our group has previously conducted an extensive research on biological properties of organoruthenium(II) pyrithionato complexes,^35–38^ however, zinc appears to be more suitable to combat COVID-19. In addition to antiviral properties, zinc pyrithione has been thoroughly investigated before for other antimicrobial properties and is now used in commercial anti-dandruff shampoos,^32^ and as antifouling agent in paints.^39^

In this study, a small library of zinc compounds containing the ligand pyrithione and its analogues (**1a–h**) was prepared and tested on two SARS-CoV-2 targets, cathepsin L and PL^Pro^. In addition, the ligands themselves (**a–h**) and the ruthenium complex with pyrithione **2a** were included to better understand the role of the metal centre (Fig. 1). Based on the excellent results of *in vitro* enzyme inhibition, further *ex vivo* assays were performed to investigate the inhibition of SARS-CoV-2 entry as well as its replication.

## Results

### Synthesis of pyrithiones and their zinc complexes

First, pyrithionato ligands were prepared in a two-step synthesis and then complexed with the zinc ion (Fig. 1). We prepared pyrithiones **b–e** with methyl substituents at different positions of the pyrithione backbone together with two pyrithiones **f–g** with extended aromaticity. In one of our publications, we have already shown that organoruthenium(II) chlorido and pta (1,3,5-triaza-7-phosphaadamantane) complexes with methyl-substituted pyrithione at position 3 exhibited enhanced cytotoxicity in seven cancer cell lines tested, compared with the chlorido/pta complexes with unsubstituted pyrithione.^36^ Therefore, we hypothesised that the methyl group as a donor substituent at position 3 would lead to better stability of the complexes. Based on that, we prepared another pyrithione ligand **h** with a donor group at position 3, i.e., a methoxy substituent −OCH_3_. According to the concept of hard and soft acids and bases (HSAB) concept, zinc has a borderline character. It tends to form complexes with both hard oxygen and soft sulphur donors. Pyrithiones have the ability to coordinate well to various metal ions via their sulfur and oxygen donor atoms to form metal complexes, and particularly stable complexes are formed with zinc (apparent K_d_ = 5.0 · 10^-12^ M^2^).^40^

### Crystal structures of zinc complexes

In the literature it is described that the zinc pyrithione complex **1a** crystallizes as a dimer with trigonal bipyramidal geometry, where the zinc ion is coordinated by five donor atoms (Fig. 2). Two pyrithionato units bind to the central zinc ion via *S*- and *O*-atoms to form a monomeric subunit, which is further linked by an additional Zn–O coordination bond from another monomeric subunit to form a dimer. However, in solution zinc pyrithione exists as a monomer.^42^ The crystal structure of zinc complex **1b** is comparable to that of the zinc pyrithione complex.^43^ For complex **1c**, the crystal structure was published as an ethanol solvate monomer with a distorted square-planar geometry and ligands in *trans*-configuration.^44^ However, we obtained the crystal structure of **1c** without solvent molecules, in which a distorted tetrahedral geometry around zinc was found (Supplementary Fig. 1). The crystals were obtained by liquid diffusion at room temperature using chloroform/hexane solvents. In addition, a new crystal structure for complex **1d** was obtained as a monomer with two ligands **d** coordinated to zinc (Fig. 2, Supplementary Fig. 1), forming a tetrahedral geometry similar to that of complex **1f** (Supplementary Fig. 1). Crystals for both complexes were obtained by liquid diffusion at room temperature using chloroform/heptane for **1d** and chloroform/dichloromethane solvent system for **1f.** Crystallographic data for novel crystal structures of **1c**, **1d** and **1f** are given in Supplementary Table 1. Moreover, complex **1e** is known to crystallize in a distorted tetrahedral geometry as a monomer.^45^ We have analysed the available crystal structures of zinc-pyrithione complexes as well as the structures obtained by our group (Supplementary Table 2). The complexes crystallize either as monomers or as dimers. The geometry in monomeric forms is distorted tetrahedral (τ = 0.70–0.78) and in dimeric forms distorted trigonal bipyramidal (τ = 0.51–0.59) (Supplementary Table 2).

**Fig. 2:**
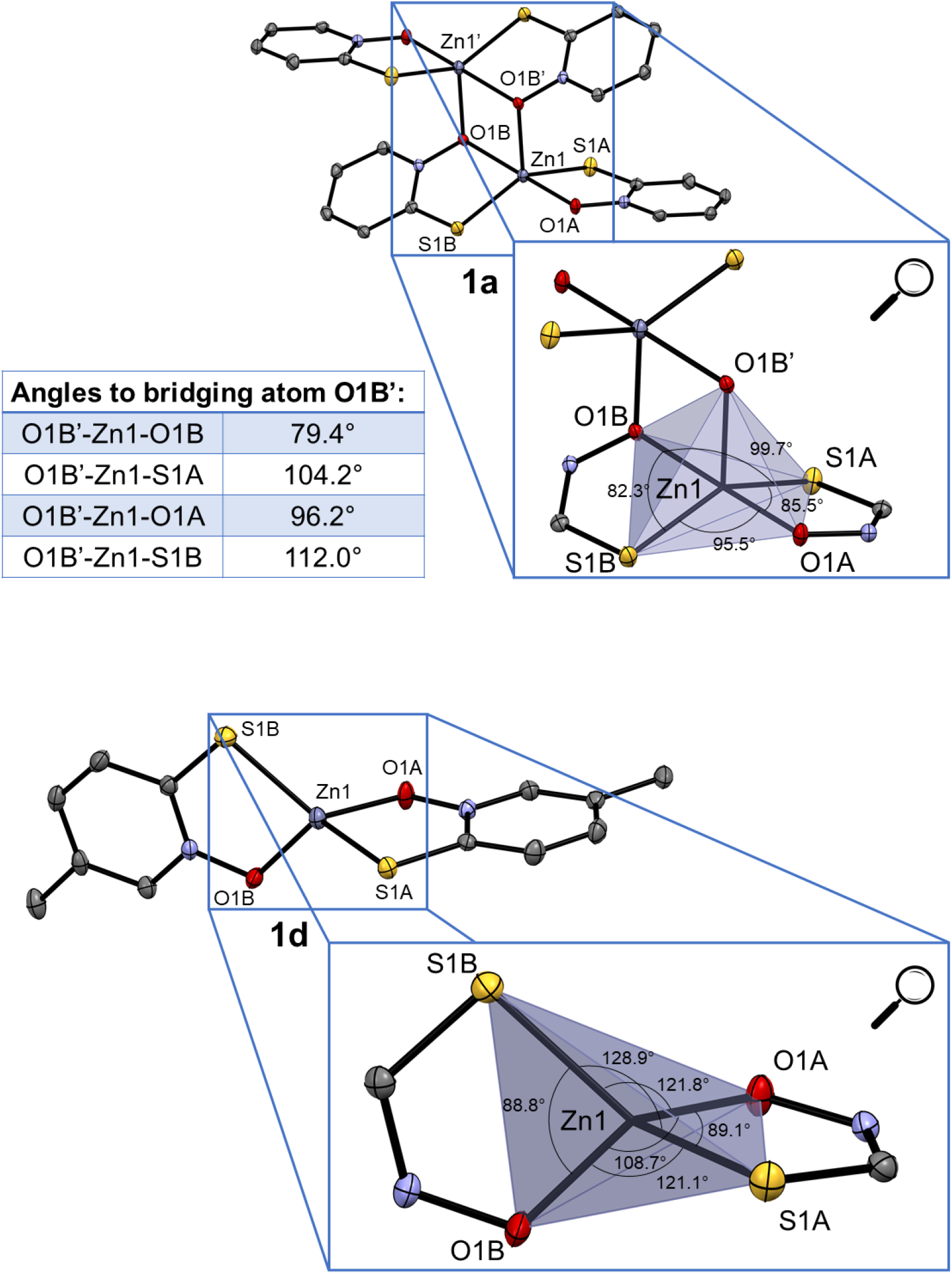
Crystal structure of complexes 1a and 1d (together with zoom of trigonal bipyramidal and tetrahedral coordination, respectively). Ellipsoids are drawn at the 35% probability level.

### UV-vis and NMR stability in biologically relevant media

UV-vis spectroscopy was first used to determine if the compounds are stable in aqueous media and therefore suitable for further biological studies (Supplementary Figs. 10–15). The same media were used as for cathepsin L and PL^Pro^ enzymatic assays, i.e. either 1% DMSO/acetate buffer (for the cathepsin L assay) or 1% DMSO/HEPES buffer (for the PL^Pro^ assay). From the UV-vis spectra, it is evident that ligand **a** and its zinc complex **1a** remain stable in acetate buffer (Supplementary Figs. 10–11). Ruthenium complex **2a** also proved to be a stable compound in this media as well (Supplementary Fig. 12). UV-vis stability measurement of the above-mentioned compounds in HEPES buffer also gave comparable curves (Supplementary Figs. 13–15). Therefore, the UV-vis stability analysis provided us with encouraging results for further biological assays.

The selected compounds (**a**, **1a**, **2a**) were further examined for their stability by ^1^H NMR spectroscopy at two selected time points appropriate for the time frame of the enzymatic assays (t = 0 and 30 min; Supplementary Figs. 16–19). The media for compounds **a** and **1a** were either i) 1% DMSO-*d*_6_/acetate buffer prepared in D_2_O (as for cathepsin L assay) or ii) 1% DMSO-*d*_6_/HEPES buffer prepared in D_2_O (as for PL^pro^ assay). For ruthenium complex **2a**, both media were used without DMSO-*d*_6_. Ligand **a** itself is a stable organic compound, showing no structural changes when tracking stability in both media over the selected time points (Supplementary Figs. 16a–b, 18a–b). Moreover, complex **1a** also proved to be stable in the media studied (Supplementary Figs. 16c–d, 18c–d). Also, ruthenium complex **2a** remains stable throughout the NMR measurements (Supplementary Figs. 17, 19). These results further supported the experimental UV-vis data and confirmed that our compounds are suitable candidates for enzymatic assays.

### Enzymatic assays

We investigated the inhibitory properties of the synthesized complexes towards two enzymes involved in the development of COVID-19, namely cathepsin L and SARS-CoV-2 PL^Pro^ (Table 1, Fig. 3, Supplementary Figs. 20–21). First, zinc complex **1a** and ruthenium complex **2a**, both of which have a pyrithione ligand bound to a metal centre, were tested to determine which metal core would be more likely to contribute to SARS-CoV-2-related target inhibition. Zinc complex **1a** inhibited both enzymes tested in the low (sub)micromolar range with IC_50_ values of 1.88 ± 0.49 μM and 0.73 ± 0.20 μM for cathepsin L and PL^Pro^, respectively. In contrast, ruthenium complex **2a** with the same pyrithione ligand showed less effective inhibition in the micromolar range with IC_50_ values of 116 ± 23 μM for cathepsin L and 14.52 ± 2.49 μM for PL^Pro^. Therefore, we decided to further investigate the inhibitory potential of analogues of zinc complex **1a**, i.e., zinc complexes with methyl-substituted pyrithiones at different positions **1b–e** and zinc complex with methoxy-substituted pyrithione **1h** as well as zinc complexes with pyrithiones with an extended aromatic backbone **1f–g**.

**Table 1:** Determined IC_50_ values for zinc and ruthenium complexes towards cathepsin L and SARS-CoV-2 PL^Pro^. Data are presented as mean ± s.e.m. (N = 3).

| **Complex** | **IC<sub>50</sub> [μM]** |  |
| --- | --- | --- |
|  | **Cathepsin L** | **PL<sup>Pro</sup>** |
| **1a** | 1.88 ± 0.49 | 0.73 ± 0.20 |
| **1b** | 0.35 ± 0.12 | 1.09 ± 0.62 |
| **1c** | 0.41 ± 0.08 | 0.54 ± 0.25 |
| **1d** | 0.44 ± 0.12 | 0.22 ± 0.08 |
| **1e** | 0.24 ± 0.05 | 0.26 ± 0.18 |
| **1f** | 0.14 ± 0.05 | 0.26 ± 0.10 |
| **1g** | NA | 1.66 ± 0.31 |
| **1h** | 0.74 ± 0.13 | 0.14 ± 0.05 |
| **2a** | 116 ± 23 | 14.52 ± 2.49 |

**Fig. 3:**
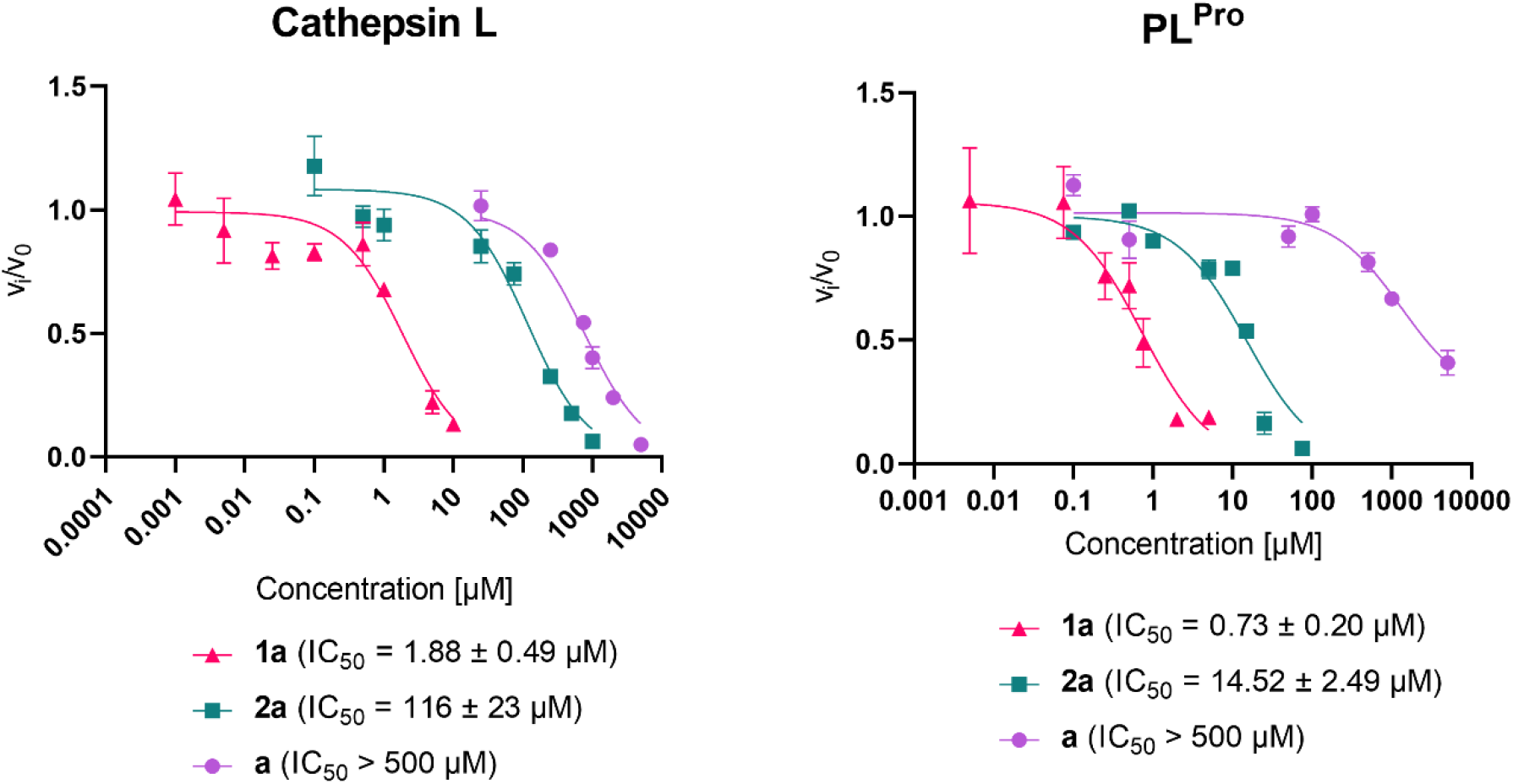
Effect of pyrithione complex with zinc 1a and with ruthenium 2a on cathepsin L and SARS-CoV-2 PL^Pro^. The effect of pyrithione (ligand a) is also shown. Data are presented as mean ± s.e.m. (N = 3).

The introduction of a methyl group at positions 3 (**1b**), 4 (**1c**), 5 (**1d**) and 6 (**1e**) resulted in somehow stronger inhibition of cathepsin L, compared with the unsubstituted complex **1a** (IC_50_ = 1.88 ± 0.49 μM), with an IC_50_ of 0.35 ± 0.12 μM for **1b**, 0.41 ± 0.08 μM for **1c**, 0.44 ± 0.12 μM for **1d**, and 0.24 ± 0.05 μM for **1e**. Similar values were obtained for the complex with isoquinoline ligand (**1f**, IC_50_ = 0.14 ± 0.05 μM) and the complex with methoxy-substituted ligand (**1h**, IC_50_ = 0.74 ± 0.13 μM). Interestingly, the zinc complex with quinoline-derived pyrithione **1g** showed complex behaviour with cathepsin L that prevented IC_50_ calculation. In addition, it is important to note that the ligands themselves are much weaker cathepsin L inhibitors compared to their zinc complexes (Supplementary Table 3).

A reverse trend was observed for PL^Pro^ activity in the case of zinc complex **1b** with methyl-substituted pyrithione at position 3, which slightly decreased the inhibitory activity of complex **1b** (IC_50_ = 1.09 ± 0.62 μM) compared to **1a** (IC_50_ = 0.73 ± 0.20 μM). On the other hand, the methyl substituents at the positions 5 and 6 contributes to increased inhibitory strength of complex **1d** and **1e** (IC_50_ = 0.22 ± 0.08 μM and 0.26 ± 0.18 μM, respectively), similar to complex **1f** with isoquinoline substituent (IC_50_ = 0.26 ± 0.10 μM) and complex **1h** with methoxy substituent (IC_50_ = 0.14 ± 0.05 μM). However, it is important to note that complex **1f** cannot fully inhibit the enzyme. Complex **1c** inhibit PL^Pro^ in a similar manner as complex **1a** with IC_50_ value of 0.54 ± 0.25 μM and complex **1g** is a slightly weaker inhibitor (IC_50_ = 1.66 ± 0.31 μM). In addition, it is important to note that the ligands themselves are again much weaker PL^Pro^ inhibitors compared to their zinc complexes (Supplementary Table 3). Our study has shown that substitution of parent pyrithione compound with selected groups has some influence on the biological response (enzyme inhibition) but none of the substituted compounds was much more effective. However, we believe that introduction of other groups might bring greater changes and should be tested in the future.

### *Ex vivo* SARS-CoV-2 entry and replication-competent assays

Following the promising results of the enzymatic assays, we also wanted to test the efficacy of the selected compounds in an *ex vivo* physiological system of SARS-CoV-2 entry and infection, recently developed and derived from primary human lung tissue (HLT).^46^ Although some differences in the inhibition of cathepsin L and PL^Pro^ were observed for zinc complexes **1b–h** with various pyrithione derivatives, the differences were not significant compared to **1a**. Since the antimicrobial properties of zinc pyrithione **1a** have been well studied and this compound is also present in commercial shampoos, we decided to test complex **1a** in an *ex vivo* system for potential antiviral properties. In addition, ligand **a** was tested as a control and ruthenium complex **2a** was tested to investigate the importance of the metal core of the complex in inhibiting viral entry and replication.

The viral entry assay was used to determine how effective the compounds tested (**a**, **1a**, **2a**) were in preventing the entry of pseudoviral particles carrying the SARS-CoV-2 S protein into HLT cells at concentrations of 30 μM, 10 μM, 1 μM, and 0.5 μM (Fig. 4a). Viral entry was inhibited in a concentration-dependent manner. At 30 μM, zinc and ruthenium complexes, **1a** and **2a**, respectively, had comparable effects with an inhibition of 93.12% and 93.16% respectively, while ligand **a** showed an inhibition of 86.41%. The observed difference in inhibition between the complexes is more pronounced at 10 μM, where zinc complex **1a** inhibited viral penetration by 91.18%, ruthenium complex **2a** by 83.9% and ligand **a** by 74.34%. When ligand **a** was tested at 1 μM and 0.5 μM, it inhibited viral entry by 16.67% and 15.33% respectively, while ruthenium complex **2a** showed an inhibition of 36.7% and 34.54% respectively, and zinc complex **1a** showed 51.80% and 40.44% inhibition at the same concentrations. Thus, calculated IC_50_ values were 0.84 μM for **1a**, 4.66 μM for **2a** and 5.19 μM for **a**. Similar to enzyme inhibition assays, ligand **a** was therefore found to be the worst inhibitor of viral entry, followed by ruthenium complex **2a** and zinc complex **1a**. Moreover, zinc also contributes to better inhibition of viral entry than ruthenium.

**Fig. 4:**
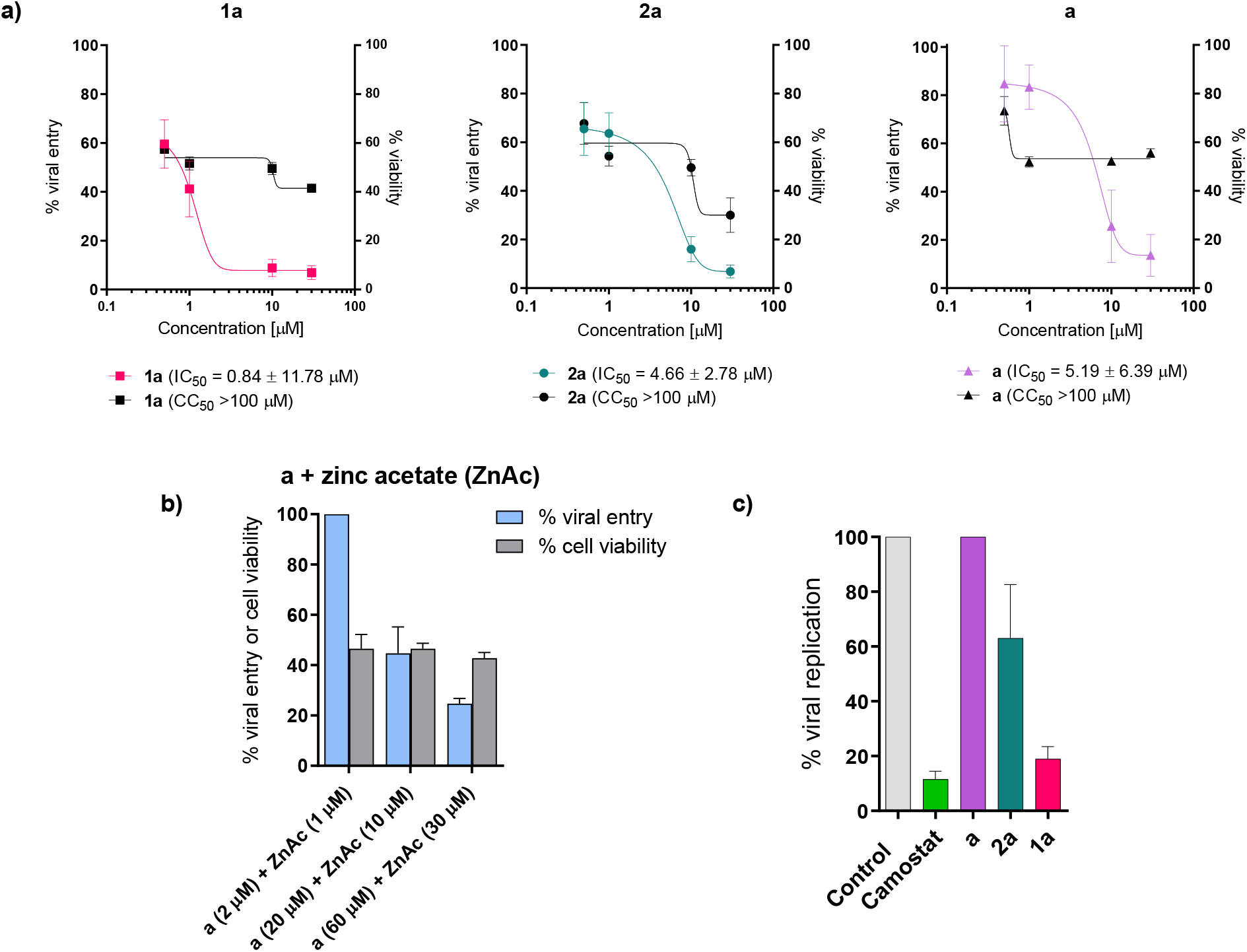
Viral entry, viral replication, and cell survival assays. a) Viral entry assay of **a**, **1a** and **2a** together with cell viability. Data are presented as mean ± s.e.m (n=3 independent experiments with technical triplicates for viral entry and duplicates for cell viability). b) Viral entry assay after the application of pyrithione **a** and zinc acetate together with cell viability. Data are presented as mean ± s.e.m (n=1 with technical triplicates for viral entry and duplicates for cell viability). c) Viral replication-competent assay for **a**, **1a** and **2a**. Data are presented as mean ± s.e.m (n=1 with technical triplicates). Camostat was used as a positive control of viral inhibition.

Besides, viral entry inhibition potential of the mixture of zinc acetate and pyrithione **a** was tested (molar ratio 1: 2, c(zinc acetate) = 1 μM, 10 μM and 30 μM; Fig. 4b). In all cases, zinc pyrithione **1a** was found to inhibit SARS-CoV-2 entry better than the above combination at all concentrations tested.

In addition to viral entry inhibition, the viability of HLT must also be taken into consideration. As expected, highest cell survival was observed at lowest tested concentration (0.5 μM) where cell survival reached 72.91%, 67.70% and 57.39% for **a**, **2a** and **1a**, respectively. Ligand pyrithione **a** at concentration 1–30 μM showed comparable cell viability of 51.91–55.51%. In the case of ruthenium complex **2a** at 30, 10, and 1 μM, cell viability was concentration-dependent, resulting in 30.02–54.26% of viable cells, while for zinc complex **1a** cell viability was 41.45–51.64%. Interestingly, cell viability remained constant (42.77-46-53%) when a combination of zinc acetate and pyrithione was tested at various concentrations.

Results for viral replication-competent assay are shown in Fig. 4c. Zinc complex **1a** inhibited 81.03% of SARS-CoV-2 replication at 30 μM and ruthenium complex **2a** prevented only 36.96% of viral replication. In addition, ligand **a** showed no effect on viral replication.

## Discussion

During the pandemic, much attention was paid to zinc because of its antiviral properties. Zinc improves mucociliary clearance (removal of viral particles from the lungs) and strengthens the integrity of the respiratory epithelium. Importantly, increased Zn^2+^ concentrations may also improve antiviral immunity by upregulating IFNα and prevent inflammation by inhibiting NF-κB signalling. In addition, hypozincemia is associated with loss of smell and taste^10–12,17^ and is reported to lead to higher severity and mortality of COVID-19 patients.^47^

As mentioned in the introduction, Zn^2+^ can indirectly inhibit SARS-CoV-2 targets such as ACE2 receptors, furin, RdRp, and M^Pro^. However, it is important to address the issue of zinc bioavailability. A randomized clinical trial suggests that oral treatment with zinc gluconate alone or in combination with ascorbic acid does not significantly shorten the duration of the COVID-19 symptoms,^48^ whereas zinc sulphate tablets may improve survival in COVID-19 patients. However, the results are not clinically applicable due to various limitations of the study.^49^ On the other hand, an ongoing clinical trial of intravenous administration of ZnCl_2_ has so far confirmed its safe use and shown an increase in serum zinc levels.^50,51^ However, oral administration of the drug is much more patient-friendly and is preferred over intravenous use. Ionophores, molecules that enable the transport of metal ions across a lipid cell membrane, increase zinc uptake and facilitate zinc absorption, thus combating COVID-19.^13,52–54^ In a multi-center study testing zinc sulphate and hydroxychloroquine as potential treatments, increased hospital discharge and decreased in-hospital mortality were observed.^55^ However, hydroxychloroquine as a therapeutic against COVID-19 should be considered with great caution as it may significantly increase the risk of cardiac arrhythmias. Similarly, a high risk of cardiac dysfunction has been reported for chloroquine.^56–58^ Therefore, the choice of an appropriate chelating agent for zinc is of utmost importance.

Pyrithione **a** is a known ionophore with the ability to bind zinc ions, increase their intracellular concentration^59^ and as such has recognized antiviral activity against Coronavirus (SARS-CoV), Arterivirus, Coxsackievirus, Mengovirus, Picornavirus and Rhinovirus.^60^ A decade ago, te Velthius *et al*. demonstrated that the combination of Zn^2+^ ions and pyrithione **a** effectively inhibited the replication of SARS-CoV via Zn^2+^ suppression of RdRp.^24^ In addition, zinc pyrithione **1a** is an established antimicrobial agent used in commercial shampoos for dandruff^32^ treatment and exerts antifungal activity by damaging iron-sulfur clusters.^31–33^ Recently, Maio *et al*. have suggested that Fe-S clusters act as cofactors of the SARS-CoV-2 RdRp enzyme and therefore represent another potential target to combat COVID-19.^34^ Therefore, pyrithione **a** and its zinc complex **1a** have attracted much attention to combat SARS-CoV-2 during the COVID-19 pandemic because of the aforementioned antimicrobial effects. However, no systematic study on zinc-pyrithione complexes was reported so far.

In the present study, we focused on two previously less studied but still very important targets of SARS-CoV-2, namely cathepsin L, which is involved in viral entry, and the protease PL^Pro^, which is involved in viral replication. Cathepsin L is a cysteine protease that cleaves the spike protein once SARS-CoV-2 is in the endolysosome within the host cell, and as such allows internalization of the virus in a manner other than TMPRSS2 and furin. Cathepsin L has been reported to be overexpressed after infection with SARS-CoV-2, allowing further progression of infection and leading to a vicious cycle: the more cathepsin L is expressed, the more severe the progression of COVID-19. Therefore, cathepsin L is a promising target to prevent SARS-CoV-2 internalization and further replication.^61^ In this study, we demonstrated that zinc pyrithione **1a** is a promising cathepsin L inhibitor with an IC_50_ value in the micromolar range (1.88 ± 0.49 μM). Although some differences in cathepsin L inhibitory activity were observed between the tested analogous zinc complexes **1b–h**, the methyl/methoxy substituents on pyrithione and its scaffold extension did not have a major influence. On the other hand, the binding of the pyrithione ligands **a–h** to the zinc core contributes significantly to the inhibition of cathepsin L and PL^Pro^ since the ligands themselves are only weak inhibitors (Supplementary Table 3). Remarkably, ruthenium pyrithione complex **2a** was also a less potent cathepsin L inhibitor (IC_50_ = 116 ± 23 μM), indicating the importance of metal core selection. Similar behaviour of the tested compounds was observed for the cysteine protease PL^Pro^, with zinc complexes being the most efficient PL^Pro^ inhibitors, and all exhibiting IC_50_ values in the submicromolar range (e.g IC_50_(**1a**) = 0.73 ± 0.20 μM), followed by ruthenium complex **2a** (IC_50_ = 14.52 ± 2.49 μM) and ligands (IC_50_ not determined). Similar to our results of PL^Pro^ inhibition, zinc pyrithione **1a** also effectively inhibited both proteases from SARS-CoV, i.e. PL^Pro^ with an IC_50_ value of 3.7 μM^27^ and M^Pro^ with a Ki value of 0.17 μM.^25,26^ In addition, zinc pyrithione **1a** also inhibited SARS-CoV-2 M^Pro^ with IC_50_ values in the submicromolar range (IC_50_ = 0.9687 μM^28^ and 0.1 μM^29^). The promising inhibition of cathepsin L and PL^Pro^ by zinc complexes, but not by ligands alone, could be attributed to the thiophilic character of zinc^54^ and its binding to cysteine residues in the active sites of the aforementioned proteases. It is worth noting that Wang *et al*. recently reported that a mixture containing colloidal bismuth subcitrate can significantly inactivate viral cysteine proteases by binding the key cysteine residue in the active sites of PL^Pro^ and M^Pro^, thus providing a potential treatment to combat SARS-CoV-2.^62^ All encouraging data from the literature as well as our new promising findings indicate that zinc pyrithione complexes are potential zinc-containing drugs for COVID-19 oral treatment. Considering that zinc pyrithione **1a** is already approved for clinical use and its biological effects have been extensively studied, complex **1a** was selected as a promising candidate for further investigation of its anti-SARS-CoV-2 entry and replication properties in a highly physiological *ex vivo* system.^46^

The *ex vivo* system is composed of primary human lung tissue (HLT) and offers many advantages over immortalized cells derived, for instance, from kidney or colon cell lines. The HLT system provides type II alveolar cells, along with other heterogeneous cell components from lung tissue that are not subjected to any differentiation processes. Importantly, the HLT system also expresses factors that are required for SARS-CoV-2 entry, such as ACE2, CD147, TMPRSS2, and AXL. Indeed, zinc pyrithione **1a** inhibited viral entry at all concentrations tested, i.e. 0.5 μM, 1 μM, 10 μM and 30 μM, with an IC_50_ of 0.84 μM. On the other hand, pyrithione **a** and ruthenium complex **2a** showed promising inhibition only at higher concentrations tested (10 and 30 μM), whereas at 0.5–1 μM only 36.4% and 34.54% of viral entry was prevented by ruthenium complex **2a** and 16.67% and 15.33% by pyrithione **a**. Since pyrithione **a** is known to be a zinc ionophore, the combination of zinc acetate and pyrithione **a** in molar ratio 1: 2 was also tested. It was found that the combination of zinc and pyrithione produced weaker inhibition than the zinc pyrithione complexes at comparable concentrations tested. Moreover, when 1 μM zinc acetate and 2 μM pyrithione were applied to the cells, viral entry was not inhibited at all.

In addition, the *ex vivo* system used also supports the investigation of SARS-CoV-2 replication inhibition. Pyrithione **a** itself did not inhibit the replication of SARS-CoV-2. Again, zinc pyrithione **1a** was found to be the most active among the compounds tested, providing 81.03% inhibition of SARS-CoV-2 replication at 30 μM, followed by ruthenium complex **2a**, which was only able to suppress replication by 36.96%.

The biological activity of the compound is, of course, a prerequisite for its possible use as a therapeutic agent. However, the efficacy of the drug in the body depends on many parameters, including the physicochemical and ADME (absorption, distribution, metabolism, excretion) properties of the drug candidate, which influence the pharmacokinetic and pharmacodynamic properties of the potential drug. The descriptors incorporated into the SwisADME tool showed that zinc pyrithione **1a** has promising properties that are favourable for drug design (Supplementary Figure 22).^63^ Generally, the compounds with logP values in the range of 1–4 are considered to be suitable for use as an oral drug. As such predicted logPo/w value of 1.65 for complex **1a** meets this requirement. Based on the calculations, complex **1a** is expected to have high gastrointestinal absorption without permeation through the blood-brain barrier. Zinc pyrithione **1a** also meets the standards of Lipinski’s rule of five and thus represents a “drug-like” compound with favourable bioavailability. One parameter that should be considered with a little more caution is the solubility of complex **1a**, which should be taken into account when preparing a potential final peroral formulation.^63,64^ Nowadays, however, advances in technological modifications allow many innovative delivery systems to improve oral drug bioavailability, e.g., nanoformulations, liposomes, co-crystallisation, cyclodextrins, polymers etc.^65,66^

In summary, we have shown that zinc pyrithione **1a** is a promising candidate to combat COVID-19 and that coordination of two pyrithione (**a**) ligands specifically to zinc is essential for the desired antiviral activity. Although some modifications should be considered to achieve better cell viability, overall complex **1a** holds great potential in the fight against SARS-CoV-2 because of its demonstrated multistep antiviral inhibitory properties. The multi-target approach to drug design has become a common strategy to avoid the possible development of inefficient drugs against SARS-CoV-2 infections as well. No therapeutic is universally effective, so broadening the spectrum of options in the fight against COVID-19 is of paramount importance, especially due to rapidly emerging mutations of the SARS-CoV-2 virus. To date, zinc pyrithione **1a** has shown excellent inhibition of the enzyme cathepsin L, which is involved in SARS-CoV-2 entry, as well as inhibition of M^Pro^ and PL^Pro^, enzymes involved in SARS-CoV-2 replication. Moreover, *ex vivo* studies have confirmed the potential of zinc pyrithione **1a** against SARS-CoV-2 entry and replication, expanding existing knowledge of its antiviral potential. Since zinc pyrithione **1a** is already an established antimicrobial agent commercially available, repurposing this zinc-containing agent for oral at-home treatment of SARS-CoV-2 patients should facilitate and accelerate its potential clinical use.

## Methods

### General procedure for the synthesis of zinc complexes

Zinc complexes **1a–h** have been prepared according to the following modified procedure.^41^ Appropriate ligand **a–h** (2 mol. equiv.) was dissolved in methanol to which 1M NaOH aqueous solution was added dropwise to obtain pH ~ 8. Then, aqueous solution of Zn(CH_3_COO)_2_·2H_2_O (1 mol. equiv.) was added to the solution of the ligand upon which white precipitate appeared immediately. Suspension was further stirred for 1 h at room temperature. White solid was filtered off under reduced pressure and first washed with methanol to eliminate a by-product CH_3_COONa and additionally with diethyl ether. Obtained zinc complexes **1a–h** were left to dry overnight at 45 °C.

Ligands **b–g** were synthesized as previously reported.^36,38,67^ Newly prepared ligand **h** was synthesized according to the same procedure as ligands **b–e**.

Synthesis of ruthenium complex **2a** has been reported before by the Turel group.^36^

### UV-vis stability

The same buffers as for enzymatic assays were used for UV-vis stability testing (50 mM acetate buffer, pH = 5.5, and 50 mM HEPES buffer, pH = 7.4). Pyrithione **a** and zinc pyrithione **1a** were dissolved in DMSO. Mixtures of a dissolved compound in DMSO (1%) and buffer (99%) were filtered through Minisart^®^ Regenerated Cellulose Syringe Filter (25 mm, 0.2 μm). Ruthenium complex **2a** was dissolved directly in buffers. After preparation, the spectra were immediately recorded on UV-vis spectrometer with an additional recording after 30 min.

### NMR stability

Acetate and HEPES buffers were prepared in D_2_O. Pyrithione **a** and zinc pyrithione **1a** were first dissolved in DMSO-*d*_6_ to which selected buffer in D_2_O was added to obtain final 1% DMSO-*d*_6_/acetate buffer or 1% DMSO-*d*_6_/HEPES buffer solutions. Such solutions were further filtered through Minisart®Regenerated Cellulose Syringe Filter (25 mm, 0.2 μm). Ruthenium complex **2a** was dissolved in deuterated buffers only without DMSO-*d*_6_ addition and filtration. ^1^H NMR spectra were recorded immediately after solution preparations and after 30 min.

### Protein expression and isolation

Recombinant procathepsin L was prepared according to the described protocol.^68^ Recombinant PL^Pro^ was prepared according to the modified protocol.^69^ Briefly, the sequence was codon optimized, synthesized and cloned into pET28b/32 (pET28b with ampicillin resistance) using *Aco*I and *Xho*I sites. The resulting plasmid was transformed into *E. coli* strain BL21(DE3) and cells were cultured in LB medium with ampicillin (50 μg/ml) at 37 °C to an OD_600_ of 0.7. PL^Pro^ expression was induced by adding 0.5 mM IPTG and 1 mM ZnCl_2_ and cells were further grown overnight with shaking at 18 °C. They were collected by centrifugation for 10 min at 8000 g and the pellet was resuspended in 50 mM TRIS-HCl, 150 mM NaCl, 10 mM imidazole, pH 8.5. Afterwards, the cells were lysed by homogenization and sonication and the lysate was clarified by centrifugation at 30 000 g for 20 min. Supernatant was filtrated and then loaded onto a Ni-affinity column, pre-equlibrated with lysis buffer. Bound proteins were eluted in a linear imidazole gradient (10–250 mM) using chromatographic system. Fractions containing PL^Pro^ were collected and dialyzed against 20 mM TRIS, 100 mM NaCl, 2 mM DTT, pH 8.5. The protein suspension was aliquoted and stored at −80 °C.

### Enzyme assays

Autocatalytical activation of procathepsin L was performed at pH ~ 4. To 1 volume (V) of procathepsin, 5% V of 3 M acetate buffer, pH 3.8 was added and the mixture was then incubated at 37 °C. Enzyme activity was monitored using fluorogenic substrate. When maximum activity was reached, the activation was stopped by adding 10% V of a neutralizing buffer, 3 M acetate buffer, pH 5.5.

Reaction mixture for cathepsin L constituted of 50 mM acetate buffer, pH 5.5, 2 μM Z-LR-AMC, 5 μM DTT, and compound (final concentration 0 – 100 μM). PL^Pro^ reaction mixture was 50 mM HEPES buffer, pH 7.4, 50 μM Z-RLRGG-AMC, 5 μM DTT, and compound (final concentration 0 – 100 μM).

All enzymatic assays were performed at 25 °C with constant magnetic stirring. Hydrolysis of fluorogenic substrates was monitored at λ_ex_ 370 nm and λ_em_ 455 nm. Data were analysed and visualized in GraphPad Prism 9. Reaction rate *ν_z_* was defined using the following equation: 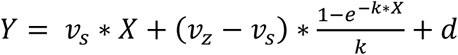, where *Y* represents measured fluorescence and *X* represents time points. Obtained reaction rates were corrected for the inner filter effect (IFE) according to the published procedure^70^. IC_50_ values were calculated using the following equation: 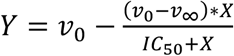, where *Y* represents IFE corrected reaction rate and *X* represents concentration of a complex. Relative reaction rates for the ligands were determined as ratio of *ν*_100 μ*M ligand*_ over *ν*_0_.

### Cell-based assays

#### Entry viral assay

Triplicates of the dilutions 30, 10, 1, and 0.5 μM of the antiviral compounds **a**, **1a**, and **2a** were tested for their antiviral effect in human lung tissue (HLT) cells using 3 different donors. Pyrithione **a** was also tested in combination with zinc acetate dihydrate (ZnAc) at different concentrations (2, 20, and 60 μM of pyrithione **a**), but maintaining a ratio of 2:1. Drug dilutions were freshly prepared in R10 in 96-well conic-bottom plates. HLT cells were obtained and processed as previously described,^46^ added at a density of 300,000 cells/well, and incubated with the compounds for 1h before infection. Then, MOI (multiplicity of infection) 0.1 of VSV*ΔG(Luc)-S virus harbouring the spike from the D614G variant was added to the individual wells, and plates were spinoculated at 1,200 g and 37 °C for 2 h. After infection, fresh medium containing the studied compounds was added to the wells, and cell suspensions were transferred into a new 96-well flat-bottom plate. Cells were then cultured overnight at 37 °C in a 5% CO_2_ incubator. Each plate contained the following controls: no cells (background control), cells treated with medium (mock infection), cells infected but untreated (infection control), and cells infected and treated with 30 μM of camostat (positive drug control). In parallel, duplicates of the tested dilutions in each assay were run to monitor drug cytotoxicity by luminescence, as previously described.^46^ After 20h, cells were incubated with Britelite plus reagent (Britelite plus kit; PerkinElmer) and then transferred to an opaque black plate. Luminescence was immediately recorded by a luminescence plate reader (LUMIstar Omega). To evaluate cytotoxicity, we used the CellTiter-Glo Luminescent kit (Promega), following the manufacturer’s instructions. Data was normalized to the mock-infected control, after which IC_50_ and CC50 values were calculated using Graph-Pad Prism 7.

#### Drug validation with replication-competent SARS-CoV-2

These experiments were performed in BSL3 facilities (Centre de Medicina Comparativa i Bioimatge de Catalunya (CMCiB) in Badalona, Spain. HLT tissue blocks were cultured at a density of 25-35 blocks/well in a 24-well plate and incubated with compounds **a**, **1a**, and **2a** at 30 μM for at least 1h before infection. Then, tissue blocks were infected with a MOI 0.5 of the SARS-CoV-2 viral isolate, and the plate was incubated for 3h at 37 °C and 5% CO_2_. After infection, samples were extensively washed with PBS 1X to eliminate residual virus and transferred into a new 24-well plate with fresh media containing the antiviral drugs. 24h post infection, 100 μl of supernatant was collected in tubes containing 100 μl of DNA/RNA Shield (ZymoResearch) for SARS-CoV-2 inactivation. For each experiment, a negative control, cells treated with only medium, and a positive control, cells incubated in the presence of the virus alone, were included. Percentage of viral infection was calculated by RT-PCR, as previously described.^46^

## Data availability

The authors declare that the main data supporting the findings of this study are available within the paper and its supplementary information files. Any additional data can be obtained from the corresponding authors MJ.B. and I.T. upon request.

## Acknowledgements

We would like to acknowledge EATRIS Academic Collaboration and their COVID-19 Fast Response Service for the coordination of new collaboration establishment between research group of prof. dr. I. Turel and dr. María J. Buzón. We would also like to thank Maša Masič for the help with the synthesis and Tjaša Rijavec for the crystallization of complex **1d**. The authors acknowledge the financial support from the Slovenian Research Agency (research core funding No. P1-0175 together with an increase in research programme funding related to the COVID-19 pandemic), and the grant from the Health Department of the Government of Catalonia (DGRIS 1_5) and the Fundació La Marató TV3 (grants 202104FMTV3 and 202112FMTV3, to M.G. and MJ.B. MJ.B is supported by the Miguel Servet program funded by the Spanish Health Institute Carlos III (CP17/00179).

## Author contributions

MJ.B, M.N. and I.T. supervised the project. J.Kla. A.D., MJ.B. and I.T. conceived and planned the experiments. J.Kla. synthesized and characterized the compounds. J.Klj. performed X-ray analysis and interpreted the data. A.D. and J.Kla. carried out UV-vis and NMR stability experiments. A.D. and M.N. performed cloning, protein expression and purification. A.D performed enzymatic assays. D.P performed viral entry. JG.E performed the replication-competent virus assays. M.G supervised the recollection of the lung tissues, and MJ.B supervised all the *ex vivo* work. J.Kla. and A.D. contributed to original draft preparation of manuscript. J.Klj., M.N., MJ.B., I.T. contributed to reviewing and editing of the manuscript. I.T. and MJ.B. provided funding acquisition. All authors have read and agreed to the published version of the manuscript.

