## Supplementary Information for "Zinc pyrithione is a potent inhibitor of PL^Pro^ and cathepsin L enzymes with *ex vivo* inhibition of SARS-CoV-2 entry and replication"

### 1. General information

Ligand pyrithione **a**, starting materials and other reagents as well as solvents for the synthesis of ligands **b**–**h**, complexes **1a**–**h** and **2a**, were purchased from commercial suppliers (Fluorochem, Alfa Aesar, Riedel-de-Haën, Merck) and used as received without further purification.

The progress of the reaction was monitored by thin layer chromatography using pre-coated TLC sheets ALUGRAM® SIL G/UV254 (Macherey–Nagel) visualised with UV lamp (254 nm). The column chromatography of the ligands was carried out on Merck silica gel 60 (35–70 mm). $^{1}$H NMR spectra for compound characterization were acquired on Bruker Avance III 500 spectrometer at room temperature at 500 MHz. $^{1}$H NMR stability spectra were obtained on NMR Bruker AvanceNeo 600 MHz spectrometer at room temperature at 600 MHz. All chemical shifts (δ) in the spectra of the complexes are referenced to the residual peaks of deuterated solvent $(CD_3)_2SO$, $CDCl_3$ and $D_2O$ at 2.50 (referenced to the central line of a quintet), 7.26 and 4.79 ppm, respectively and are reported in ppm. Coupling constants (*J*) are reported in Hz. Spectra were processed in MestReNova 11.0.4. The multiplicity of the signals is abbreviated as s – singlet, d – doublet, t – triplet and m – multiplet. IR spectra were obtained on a Bruker FTIR Alpha Platinum ATR spectrometer. High resolution mass spectrometry was performed on an Agilent 6224 Accurate Mass TOF LC/MS system. Elemental analysis (C, H, N) was carried out on a PerkinElmer 2400 II instrument. UV-vis spectroscopy was performed on a PerkinElmer LAMBDA 750 UV/Vis/near-IR spectrophotometer. For X-ray structural analysis, single crystals were surrounded with silicon grease, mounted onto the tip of glass fibres and transferred to the goniometer head in the liquid nitrogen cryostream (150(2) K). Data were collected on a SuperNova diffractometer equipped with Atlas detector using CrysAlis software with monochromated Mo Kα (0.71073 Å).[1] The initial structural models were obtained via direct methods using the Olex2 graphical user interface[2] implemented in SHELXT. A full-matrix least-squares refinement on $F^2$ magnitudes with anisotropic displacement parameters for all non-hydrogen atoms using Olex2 or SHELXL-2018/3 was employed.[2,3] All non-hydrogen atoms were refined anisotropically, while the hydrogen atoms were placed at calculated positions and further treated as riding on their parent atoms. Figures depicting the structures were prepared with Mercury.[4] The crystal structures have been submitted to the CCDC and have been allocated the deposition numbers 2143703-2143706. These data are provided free of charge by The Cambridge Crystallographic Data Centre.[5]

Synthetic fluorogenic substrates benzyloxycarbonyl-Leu-Arg-7-amino-4-methylcoumarin (Z-LR-AMC; Cat. No. 4034611) and benzyloxycarbonyl-Arg-Leu-Arg-Gly-Gly-7-amino-4-methylcoumarin (Z-RLRGG-AMC; Cat. No. 4027158) were purchased from Bachem. Ni-affinity column HisTrap was purchased from GE Healthcare.

$OD_{600}$ measurements were obtained with Varian Cary 50 Bio spectrophotometer (Agilent). Protein elution was performed on ÄKTA FPLC system (GE Healthcare). Enzymatic kinetic measurements were performed on a PerkinElmer LS55 fluorescence spectrometer with PTP-1 Fluorescence Peltier System and Peltier Controlled Fluid Circulator (PCB1500).

### 2. Characterization of zinc complexes

[Zn(II)(1-hydroxypyridine-2(1*H*)-thionato)$_2$] (**1a**).

Yield: 89%. $^1$H NMR (500 MHz, $(CD_3)_2SO$)): δ = 8.43 (dd, 2H, *J* = 6.6, 1.2 Hz, Ar–*H* a), 7.61 (dd, 2H, *J* = 8.2, 1.8 Hz, Ar–*H* a), 7.28–7.22 (m, 2H, Ar–*H* a), 7.01 (td, 2H, *J* = 7.0, 1.8 Hz, Ar–*H* a) ppm. ESI-HRMS ($CH_3CN$): *m/z* calcd for $[C_{10}H_9N_2O_2S_2Zn]^+$: 316.9391; found: 316.9391. Elemental analysis calcd (%) for $C_{10}H_8N_2O_2S_2Zn$: C, 37.81; H, 2.54; N, 8.82; found: C, 37.59; H, 2.50; N, 8.59. IR selected bands (ATR): ṽ = 3102, 1457, 1199, 1147, 1087, 821, 762, 703, 582, 564 $cm^{-1}$.

[Zn(II)(1-hydroxy-3-methylpyridine-2(1*H*)-thionato)$_2$] (**1b**).

Yield: 87%. $^1$H NMR (500 MHz, $CDCl_3$): *δ* = 8.26 (dd, 2H, *J* = 6.6, 0.6 Hz, Ar–*H* b), 7.24–7.20 (m, 2H, Ar–*H* b), 6.84 (t, 2H, *J* = 7.0 Hz, Ar–*H* b), 2.52 (s, 6H, Ar–C*H*$_3$ b) ppm. ESI-HRMS ($CH_3CN$): *m/z* calcd for $[C_{12}H_{13}N_2O_2S_2Zn]^+$: 344.9704; found: 344.9702. Elemental analysis calcd (%) for $C_{12}H_{12}N_2O_2S_2Zn$: C, 41.69; H, 3.50; N, 8.10; found: C, 41.34; H, 3.40; N, 8.00. IR selected bands (ATR): ṽ = 3073, 1403, 1206, 1146, 1134, 782, 702, 630, 565 $cm^{-1}$.

[Zn(II)(1-hydroxy-4-methylpyridine-2(1*H*)-thionato)$_2$] (**1c**).

Yield: 50%. $^1$H NMR (500 MHz, $CDCl_3$): δ = 8.16 (d, 2H, *J* = 6.8 Hz, Ar–*H* c), 7.53 (dd, 2H, *J* = 1.6, 0.6 Hz, Ar–*H* c), 6.72 (dd, 2H, *J* = 6.8, 2.4 Hz, Ar–*H* c), 2.30 (s, 6H, Ar–C*H*$_3$ c) ppm. ESI-HRMS ($CH_3CN$): *m/z* calcd for $[C_{12}H_{13}N_2O_2S_2Zn]^+$: 344.9704; found: 344.9701. Elemental analysis calcd (%) for $C_{12}H_{12}N_2O_2S_2Zn$: C, 41.69; H, 3.50; N, 8.10; found: C, 41.53; H, 3.50; N, 7.97. IR selected bands (ATR): ṽ = 3072, 1614, 1469, 1197, 1138, 1085, 809, 780, 602, 488 $cm^{-1}$.

[Zn(II)(1-hydroxy-5-methylpyridine-2(1*H*)-thionato)$_2$] (**1d**).

Yield: 72%. $^1$H NMR (500 MHz, $CDCl_3$): δ = 8.14 (s, 2H, Ar–*H* d), 7.61 (d, 2H, *J* = 8.5 Hz, Ar–*H* d), 7.07 (dd, 2H, *J* = 8.5, 1.4 Hz, Ar–*H* d), 2.26 (s, 6H, Ar–C*H*$_3$ d) ppm. ESI-HRMS ($CH_3CN$): *m/z* calcd for $[C_{12}H_{13}N_2O_2S_2Zn]^+$: 344.9704; found: 344.9705. Elemental analysis calcd (%) for $C_{12}H_{12}N_2O_2S_2Zn$: C, 41.69; H, 3.50; N, 8.10; found: C, 41.39; H, 3.38; N, 8.08. IR selected bands (ATR): ṽ = 3056, 1474, 1372, 1149, 1090, 810, 739, 664, 592, 548 $cm^{-1}$.

[Zn(II)(1-hydroxy-6-methylpyridine-2(1*H*)-thionato)$_2$] (**1e**).

Yield: 54%. $^1$H NMR (500 MHz, $CDCl_3$): δ = 7.63 (dd, 2H, *J* = 8.3, 1.5 Hz, Ar–*H* e), 7.10 (t, 2H, *J* = 7.7 Hz, Ar–*H* e), 6.85 (dd, 2H, *J* = 7.7, 1.5 Hz, Ar–*H* e), 2.61 (s, 6H, Ar–C*H*$_3$ e) ppm. ESI-HRMS ($CH_3CN$): *m/z* calcd for $[C_{12}H_{13}N_2O_2S_2Zn]^+$: 344.9704; found 344.9702. Elemental analysis calcd (%) for $C_{12}H_{12}N_2O_2S_2Zn$: C, 41.69; H, 3.50; N, 8.10; found: C, 41.26; H, 3.39; N, 7.97. IR selected bands (ATR): ṽ = 3072, 1557, 1461, 1373, 1200, 1184, 889, 770, 646, 411 $cm^{-1}$.

[Zn(II)(2-hydroxyisoquinoline-1(2*H*)-thione)$_2$] (**1f**).

Yield: 93 %. $^1$H NMR (500 MHz, $CDCl_3$): δ = 8.77–8.72 (m, 2H, Ar–*H* f), 8.22 (d, 2H, *J* = 7.2 Hz, Ar–*H* f), 7.78–7.73 (m, 2H, Ar–*H* f), 7.72–7.66 (m, 4H, Ar–*H* f), 7.30 (d, 2H, *J* = 7.2 Hz, Ar–*H* f) ppm. ESI-HRMS ($CH_3CN$): *m/z* calcd for $[C_{18}H_{13}N_2O_2S_2Zn]^+$: 416.9704; found: 416.9708. Elemental analysis calcd (%) for $C_{18}H_{12}N_2O_2S_2Zn$: C, 51.75; H, 2.90; N, 6.71; found: C, 51.48; H, 2.67; N, 6.60. IR selected bands (ATR): ṽ = 3088, 2970, 1547, 1334, 1306, 1199, 948, 789, 762, 746, 667 $cm^{-1}$.

[Zn(II)(1-hydroxyquinoline-2(1*H*)-thione)$_2$] (**1g**).

Yield: 70 %. $^1$H NMR (500 MHz, $CDCl_3$): δ = 8.64 (d, 2H, *J* = 8.8 Hz, Ar–*H* g), 7.79–7.70 (m, 6H, Ar–*H* g), 7.66 (d, 2H, *J* = 8.8 Hz, Ar–*H* g), 7.53–7.48 (m, 2H, Ar–*H* g) ppm. ESI-HRMS ($CH_3CN$): *m/z* calcd for $[C_{18}H_{13}N_2O_2S_2Zn]^+$: 416.9704; found: 416.9704. Elemental analysis calcd (%) for $C_{18}H_{12}N_2O_2S_2Zn$: C, 51.75; H, 2.90; N, 6.71; found: C, 51.87; H, 2.61; N, 6.72. IR selected bands (ATR): ṽ = 2970, 1738, 1600, 1561, 1306, 902, 808, 757, 654, 512 $cm^{-1}$.

[Zn(II)(1-hydroxy-3-methoxypyridine-2(1*H*)-thionato)$_2$] (**1h**).

Yield: 57%. $^1$H NMR (500 MHz, $CDCl_3$): *δ* = 8.08 (dd, 2H, *J* = 6.5, 1.0 Hz, Ar–*H* h), 6.90–6.80 (m, 4H, Ar–*H* h), 3.98 (s, 6H, Ar–OC$H_3$ h) ppm. ESI-HRMS ($CH_3CN$): m/z calcd for $[C_{12}H_{13}N_2O_4S_2Zn]^+$: 376,9603; found: 376.9598. Elemental analysis calcd (%) for $C_{12}H_{12}N_2O_4S_2Zn$: C, 38.16; H, 3.20; N, 7.42; found: C, 37.97; H, 3.00; N, 7.47. IR selected bands (ATR): ṽ = 3108, 1553, 1441, 1424, 1292, 1222, 1067, 762, 696, 669 $cm^{-1}$.

### 3. Crystal structures

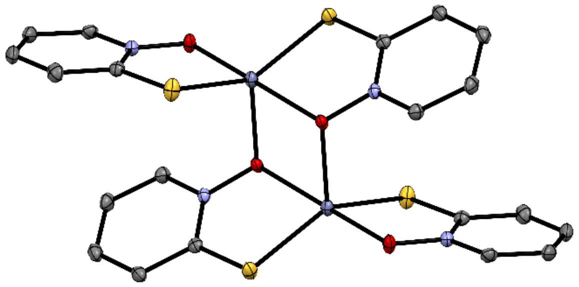

**1a***

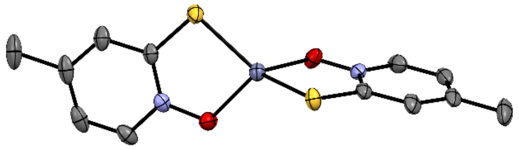

**1c**

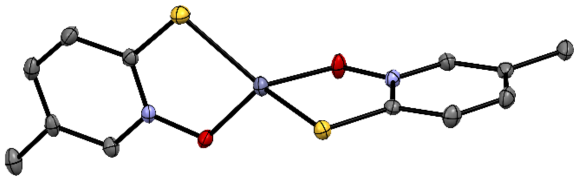

**1d**

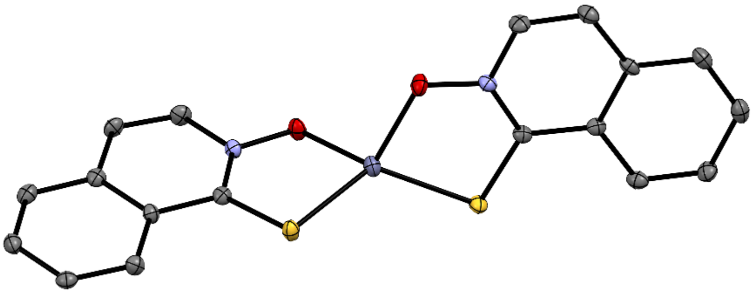

**1f**

**Supplementary Figure 1: Crystal structure of complexes 1a, 1c, 1d and 1f.** Ellipsoids are drawn at the 35% probability level. *As the structure report for **1a** dates back to 1977,[6] we decided to re-measure data on a crystal sample of this compound to obtain high-quality data.

**Supplementary Table 1: Crystallographic data for complexes 1a, 1c, 1d and 1d.**

| Compound | **1a** (moc470) | **1c** (mob156) | **1d** (mob128) | **1f** (moc457) |
|---|---|---|---|---|
| Empirical formula | $C_{10}H_8N_2O_2S_2Zn$ | $C_{12}H_{12}N_2O_2S_2Zn$ | $C_{12}H_{12}N_2O_2S_2Zn$ | $C_{18}H_{12}N_2O_2S_2Zn$ |
| Formula weight | 317.67 | 345.73 | 345.769 | 417.79 |
| Temperature/K | 150(2) | 150 | 150 | 150 |
| Crystal system | monoclinic | trigonal | monoclinic | triclinic |
| Space group | $P2_1/c$ | R-3c | C2/c | P-1 |
| a/Å | 8.3347(4) | 21.186(2) | 27.7422(12) | 7.0548(3) |
| b/Å | 10.1317(4) | 21.186(2) | 7.2234(4) | 7.0948(3) |
| c/Å | 13.6447(6) | 16.8962(9) | 13.5512(6) | 16.2539(9) |
| α/° | 90 | 90 | 90 | 99.916(4) |
| β/° | 96.748(4) | 90 | 99.630(5) | 95.707(4) |
| γ/° | 90 | 120 | 90 | 98.714(4) |
| Volume/$Å^3$ | 1144.24(9) | 6567.9(13) | 2677.3(2) | 785.53(7) |
| Z | 4 | 18 | 8 | 2 |
| $\rho_{calc}$ [g/$cm^3$] | 1.844 | 1.573 | 1.716 | 1.766 |
| μ/$mm^{-1}$ | 2.499 | 1.966 | 2.144 | 1.844 |
| F(000) | 640.0 | 3168.0 | 1413.1 | 424.0 |
| Crystal size/$mm^3$ | 0.25 × 0.25 × 0.25 | 0.7 × 0.1 × 0.1 | 1.0 × 0.9 × 0.8 | 0.1 × 0.05 × 0.03 |
| Radiation | Mo Kα (λ = 0.71073) | Mo Kα (λ = 0.71073) | Mo Kα (λ = 0.71073) | Mo Kα (λ = 0.71073) |
| 2Θ range for data collection/° | 4.922 to 54.958 | 6.35 to 54.952 | 5.84 to 54.96 | 5.132 to 54.97 |
| Index ranges | $-10 \leq h \leq 10$, $-13 \leq k \leq 13$, $-16 \leq l \leq 17$ | $-18 \leq h \leq 22$, $-27 \leq k \leq 27$, $-21 \leq l \leq 19$ | $-38 \leq h \leq 25$, $-9 \leq k \leq 10$, $-17 \leq l \leq 18$ | $-8 \leq h \leq 9$, $-9 \leq k \leq 9$, $-21 \leq l \leq 21$ |
| Reflections collected | 11634 | 5691 | 8064 | 16062 |
| Independent reflections | 2609 [$R_{int}$ = 0.0349, $R_{sigma}$ = 0.0280] | 1661 [$R_{int}$ = 0.0245, $R_{sigma}$ = 0.0207] | 3075 [$R_{int}$ = 0.0219, $R_{sigma}$ = 0.0291] | 3550 [$R_{int}$ = 0.0513, $R_{sigma}$ = 0.0428] |
| Data/restraints/parameters | 2609/0/154 | 1661/0/88 | 3075/0/174 | 3550/0/226 |
| Goodness-of-fit on $F^2$ | 1.055 | 1.251 | 1.050 | 1.137 |
| Final R indexes [I>=2σ (I)] | $R_1$ = 0.0242, $wR_2$ = 0.0534 | $R_1$ = 0.0446, $wR_2$ = 0.0923 | $R_1$ = 0.0294, $wR_2$ = 0.0738 | $R_1$ = 0.0695, $wR_2$ = 0.1942 |
| Final R indexes [all data] | $R_1$ = 0.0302, $wR_2$ = 0.0567 | $R_1$ = 0.0490, $wR_2$ = 0.0941 | $R_1$ = 0.0350, $wR_2$ = 0.0775 | $R_1$ = 0.0782, $wR_2$ = 0.1990 |
| Largest diff. peak/hole / e$Å^{-3}$ | 0.33/-0.37 | 0.43/-0.34 | 0.47/-0.34 | 2.58/-0.76 |

Comment of crystal structure of **1f:** The checkcif report contains 2 B-type alerts due to residual peaks of electron density (Q1 2.58, Q2 2.56). These peaks cannot be refined by insertion of for example solvate water molecules. The absence of solvate water molecules is confirmed by CHN elemental analysis. The anomalies are attributed to lower quality of the single crystal. Despite that, all bond lengths and angles are within the value ranges typical for zinc complexes. At present this is the best data we were able to obtain.

**Supplementary Table 2: Analysis of the available crystal structures of zinc-pyrithione complexes.**

| | | | $Zn1\text{-}O_{terminal}$ | Zn1-S | $Zn1\text{-}O_{bridge}$ | $O_{bridge}\text{-} Zn2$ | $\tau_5$ |
|---|---|---|---|---|---|---|---|
| OXPTZN[6] (Zn-pth complex, chloroform solvate) | **1a** | dimer | 2.0502 | 2.3106<br>2.315 | 2.1847 | 2.0925 | 0.59 |
| OXPZND (Zn-pth complex) | **1a** | dimer | 2.0494 | 2.3069<br>2.3075 | 2.1821 | 2.1215 | 0.52 |
| MIKZEL | **1b** | dimer | 2.049(3) | 2.306(2)<br>2.295(2) | 2.171(3) | 2.127(4) | 0.51 |
| | | | **Zn-O** | **Zn-S** | | | $\tau_4/\tau_4$' |
| TETRAL[7] (Zn-(4Me-pth) complex, 1/6 ethanol solvate) | **1c** | monomer (symmetric) | 1.968 | 2.275 | | | 0.78/0.74 |
| Mob156 (Isostructural with TETRAL, EtOH atoms unassigned) | **1c** | monomer (symmetric) | 1.973 | 2.270 | | | 0.78/0.74 |
| Mob128 (Zn-(5Me-pth) complex) | **1d** | monomer | 1.973(1)<br>1.968(1) | 2.2859(7)<br>2.2729(8) | | | 0.78/0.75 |
| CAQCEC[8] (Zn-(6Me-pth) complex) | **1e** | monomer | 1.983(2)<br>1.985(3) | 2.2636(9)<br>2.2701(8) | | | 0.76/0.70 |
| Mob457 (Zn-(N-hydroxyisoquinoline-1-thione) complex) | **1f** | monomer | 1.972(5)<br>1.965(4) | 2.289(2)<br>2.286(1) | | | 0.76/0.75 |

$\tau_5 = (\beta-\alpha)/(60°)$. For square-pyramidal geometry, the $\tau_5$ value is 0, for ideal trigonal-bipyramidal geometry, the value of $\tau_5$ is 1. $\tau_4 = [360°-(\alpha+\beta)]/(360°-2\cdot\theta)$; $\tau_4' = [(\beta-\alpha)/(360°-\theta)]+[(180°-\beta)/(360°-\theta)]$; α and β are the two greatest valence angles of coordination center, θ is tetrahedral angle 109.471°. For ideal square-planar geometry $\tau_4$ and $\tau_4'$ values are 0, for ideal tetrahedral geometry $\tau_4$ and $\tau_4'$ values are 1.

Reference for $\boldsymbol{\tau_4}$: https://pubs.rsc.org/en/content/articlelanding/2007/DT/B617136B
Reference for $\boldsymbol{\tau_4}$**'**: https://www.sciencedirect.com/science/article/pii/S0277538715000625?via%3Dihub
Reference for $\boldsymbol{\tau_5}$: https://pubs.rsc.org/en/content/articlelanding/1984/DT/DT9840001349

### 4. NMR spectra

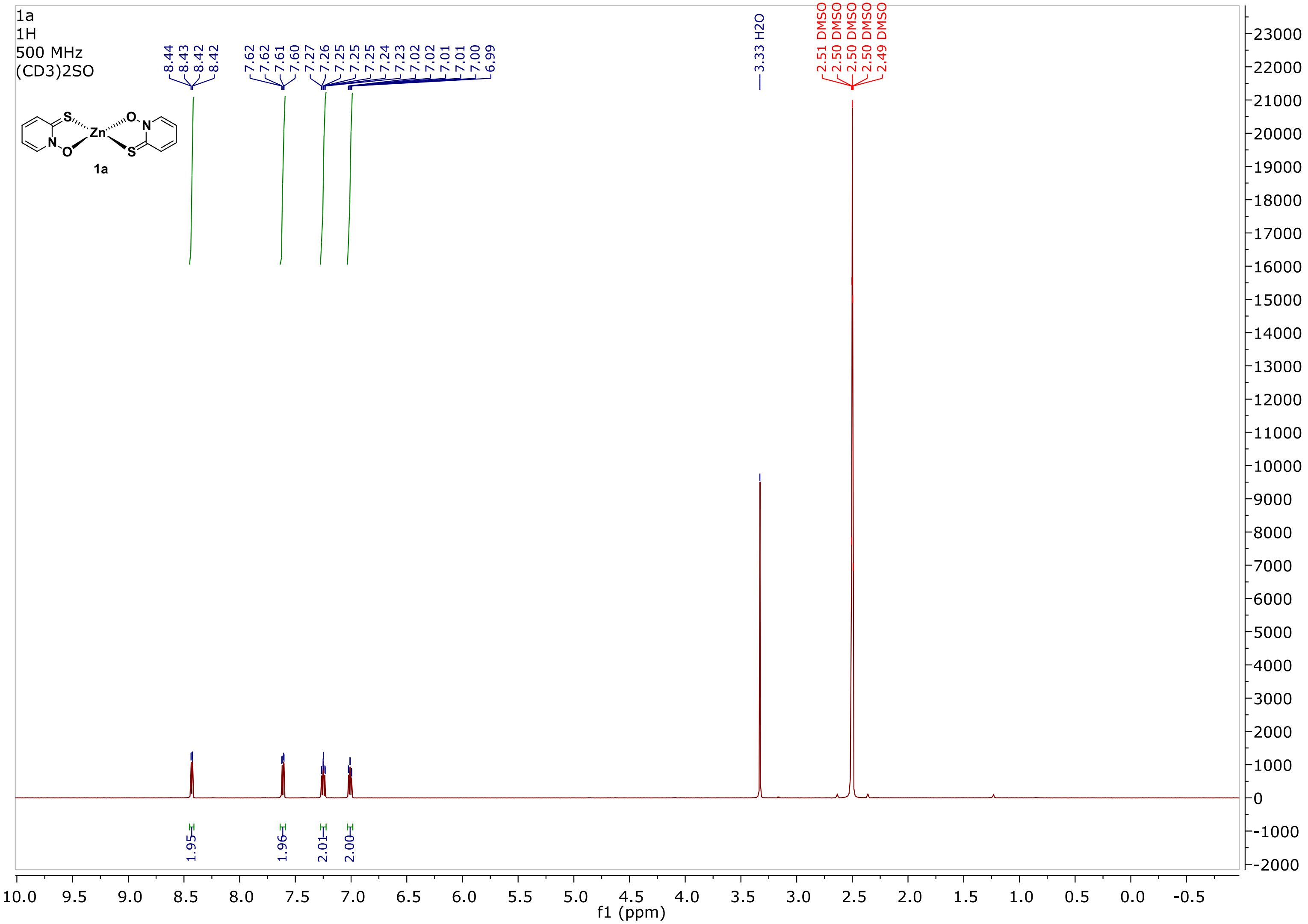

**Supplementary Figure 2: $^1$H NMR spectrum of 1a.**

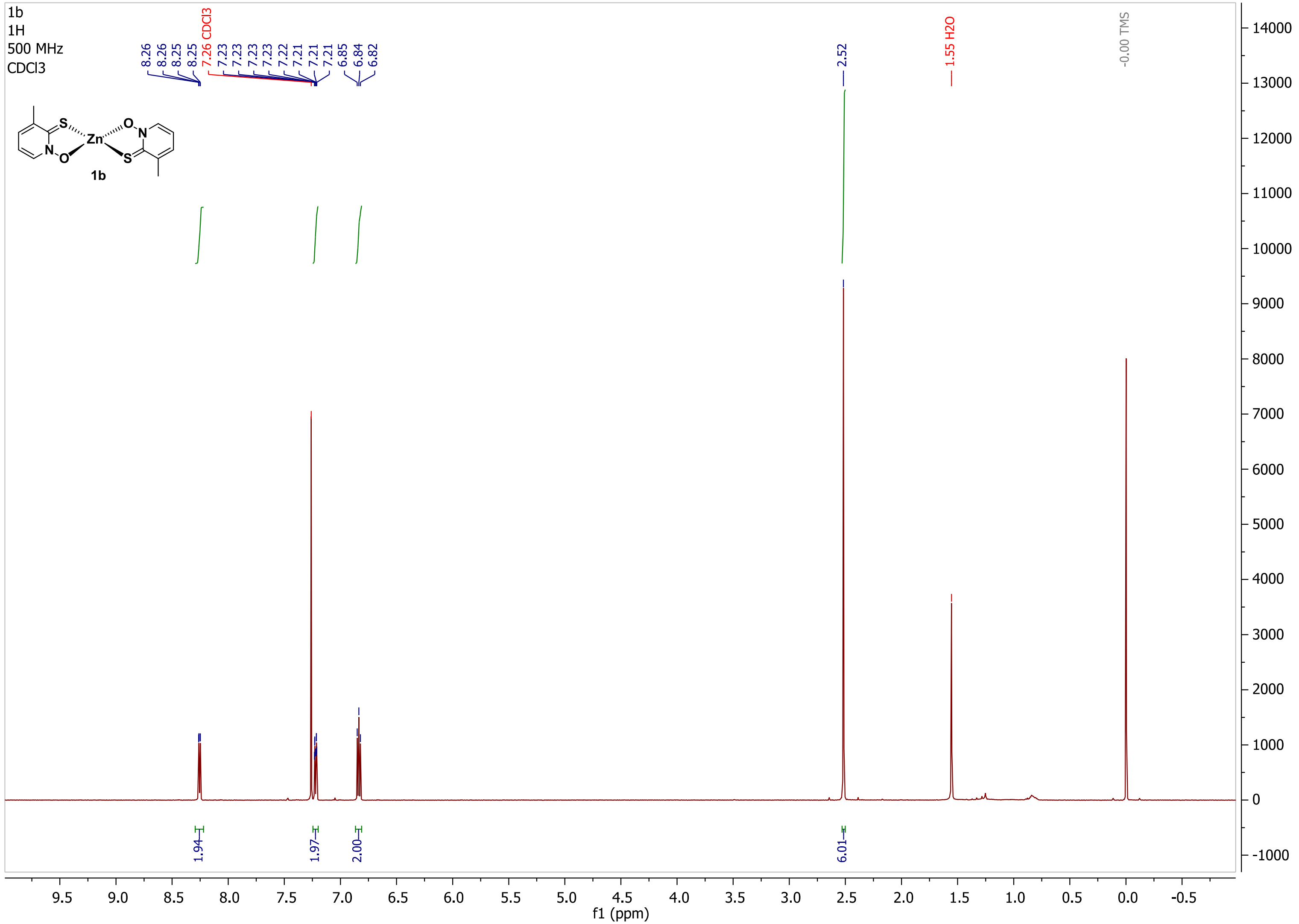

**Supplementary Figure 3: $^{1}$H NMR spectrum of 1b.**

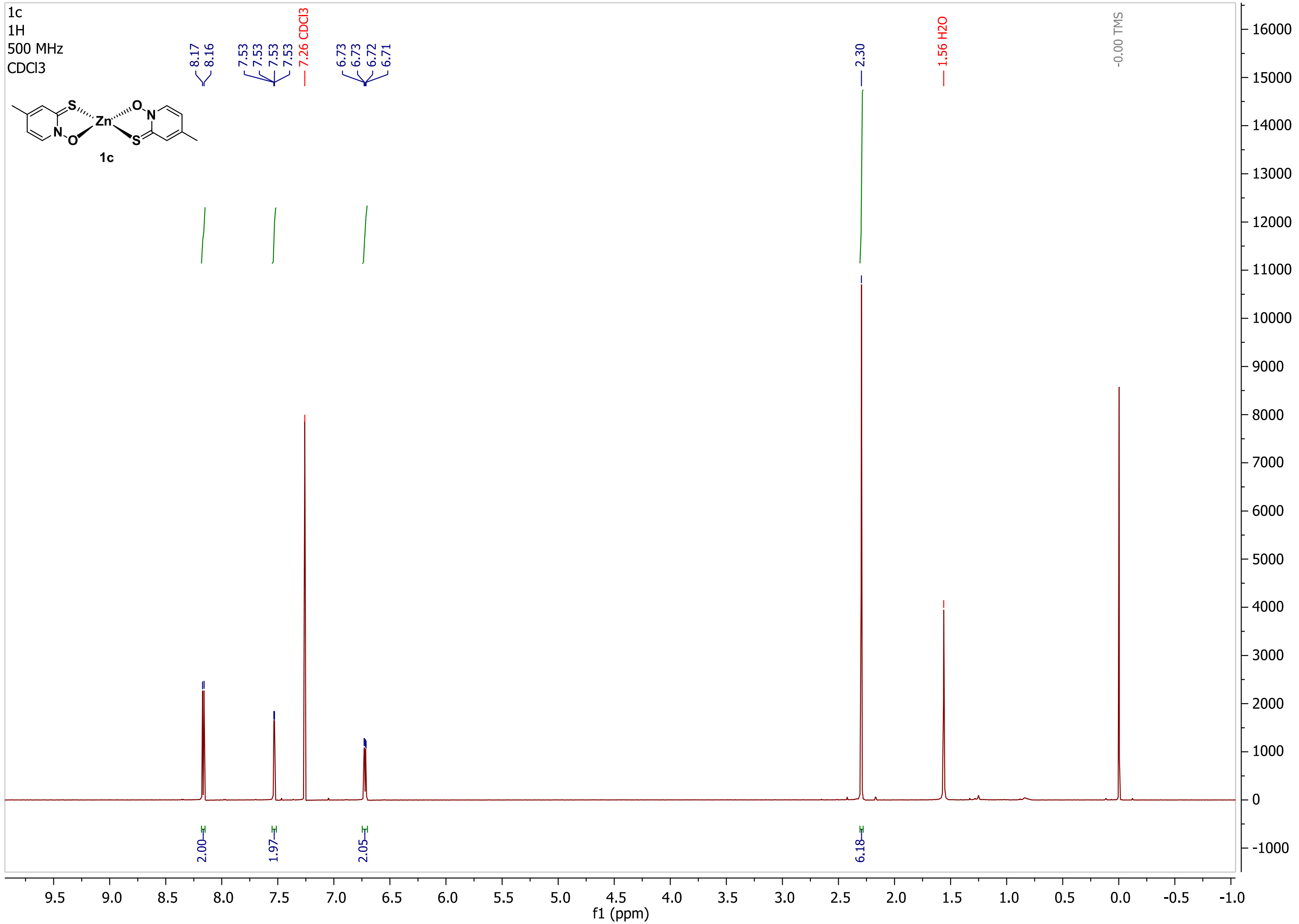

**Supplementary Figure 4: $^1$H NMR spectrum of 1c.**

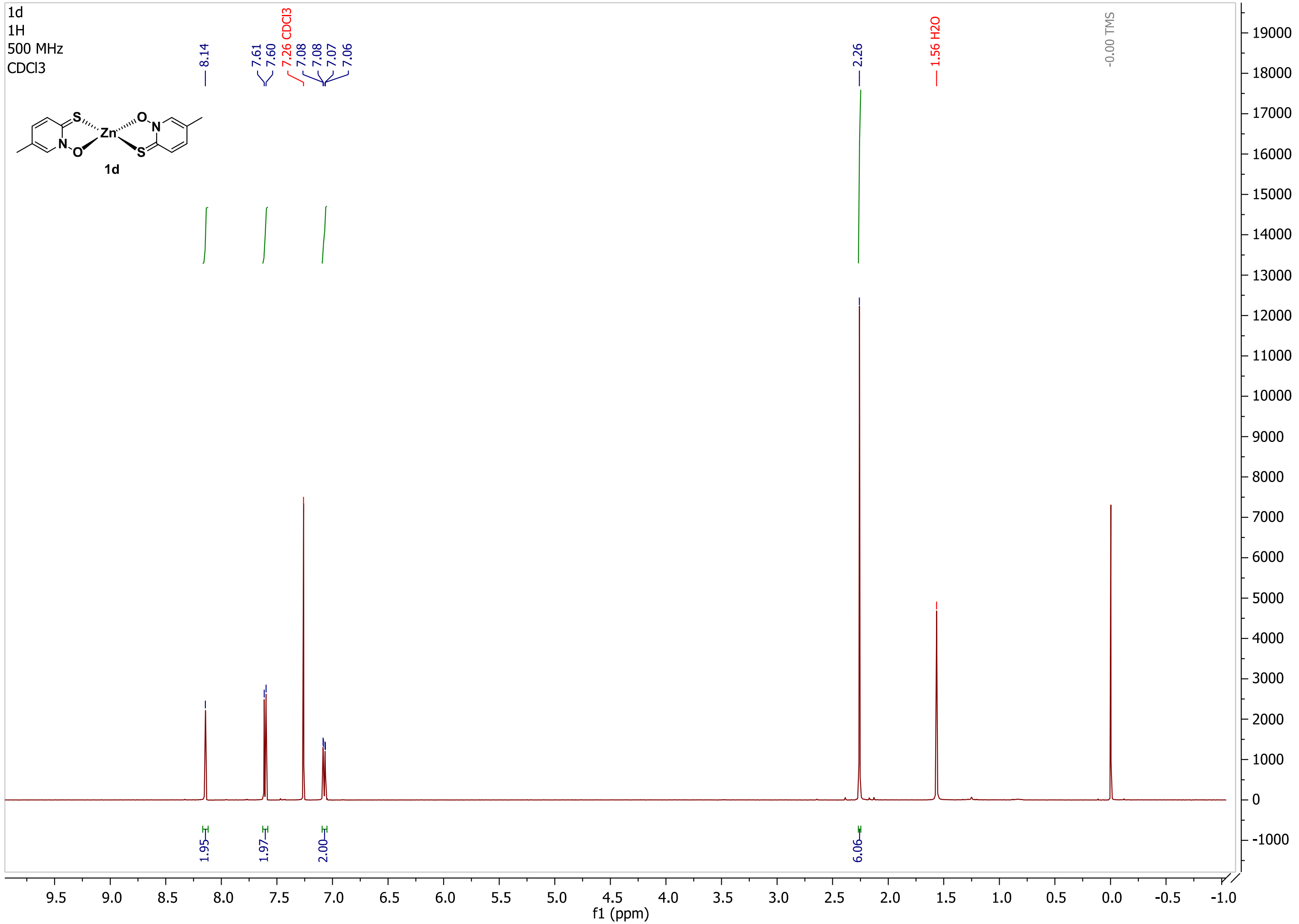

**Supplementary Figure 5: $^1$H NMR spectrum of 1d.**

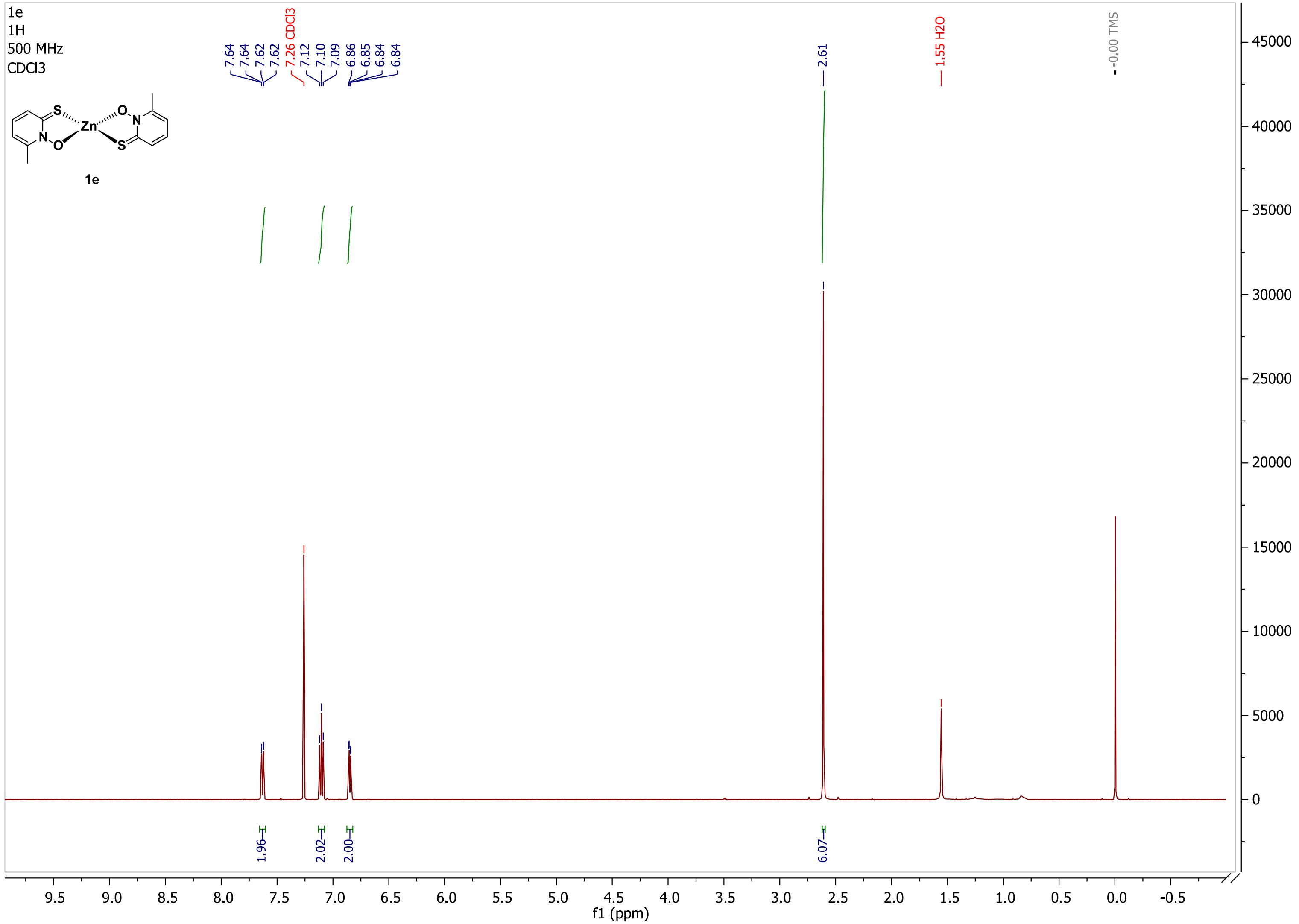

**Supplementary Figure 6: $^{1}$H NMR spectrum of 1e.**

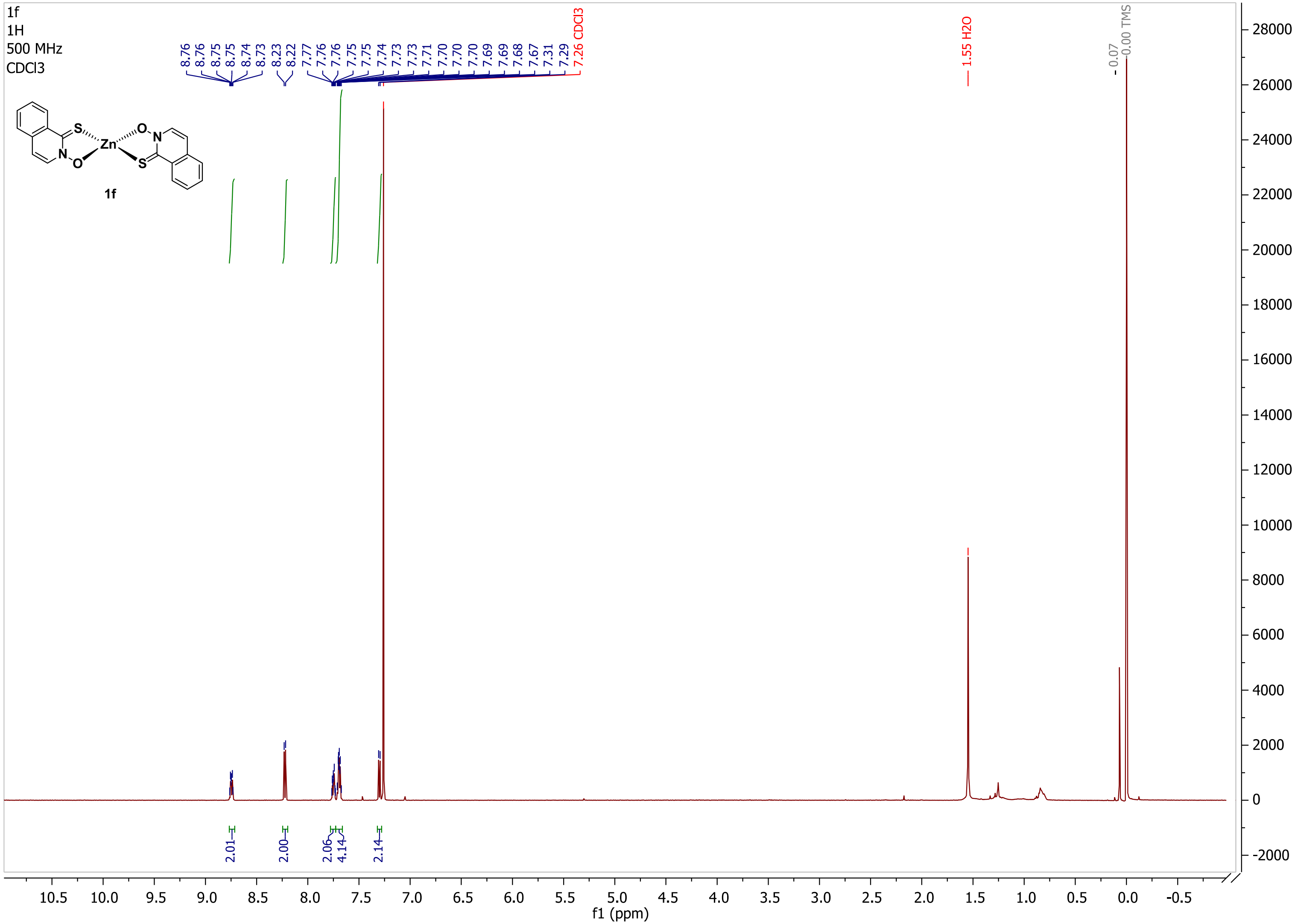

**Supplementary Figure 7: $^{1}$H NMR spectrum of 1f.**

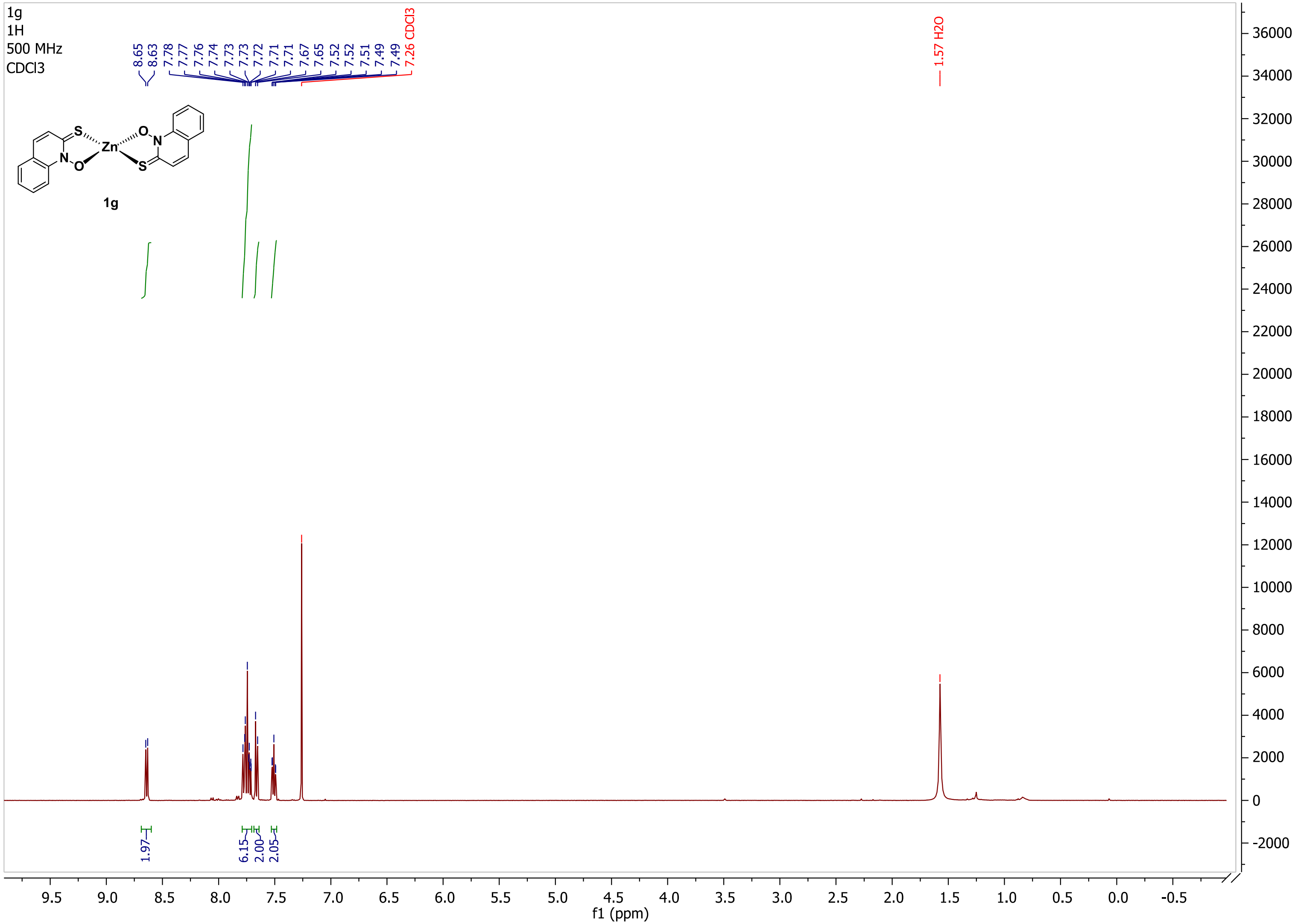

**Supplementary Figure 8: $^{1}$H NMR spectrum of 1g.**

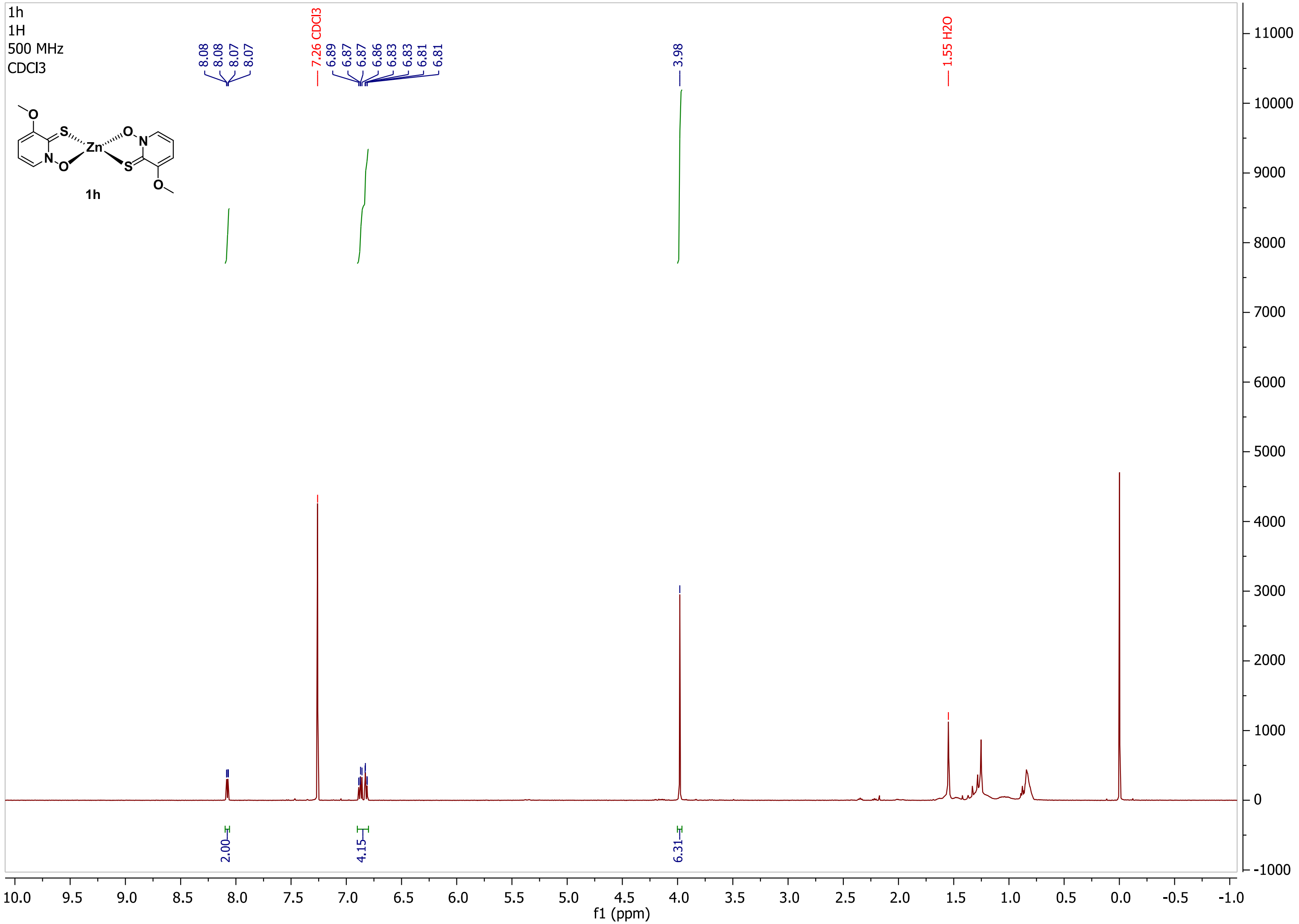

**Supplementary Figure 9: $^1$H NMR spectrum of 1h.**

### 5. UV-vis and NMR stability

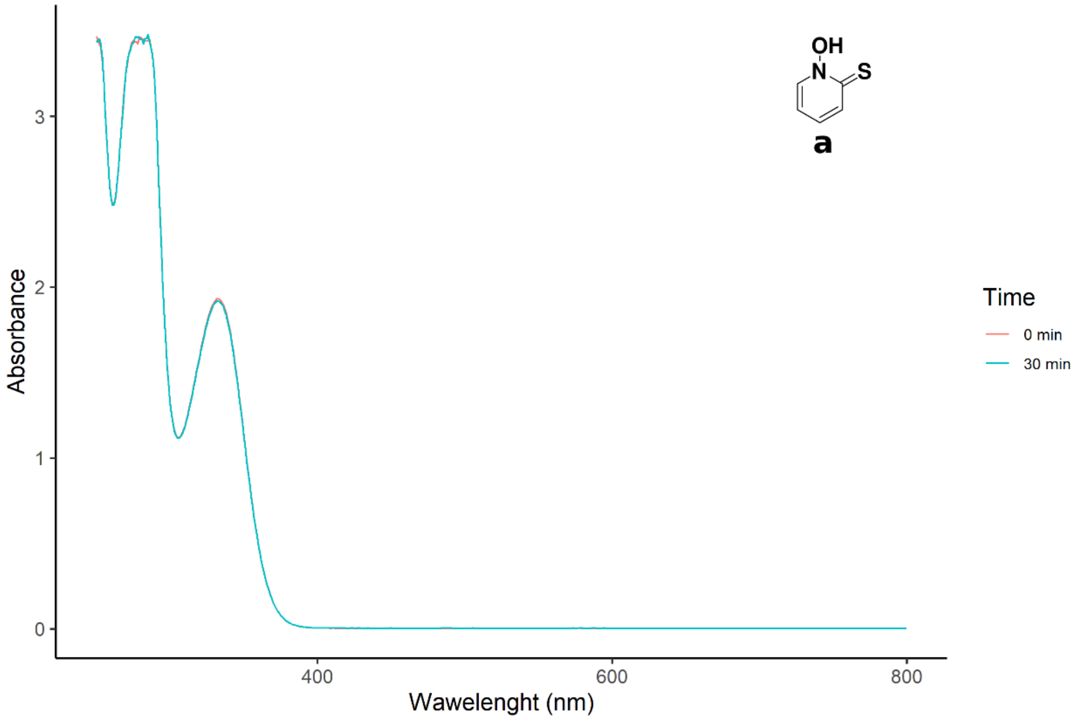

**Supplementary Figure 10: UV-vis stability spectra of ligand pyrithione a in 1% DMSO/acetate buffer.**

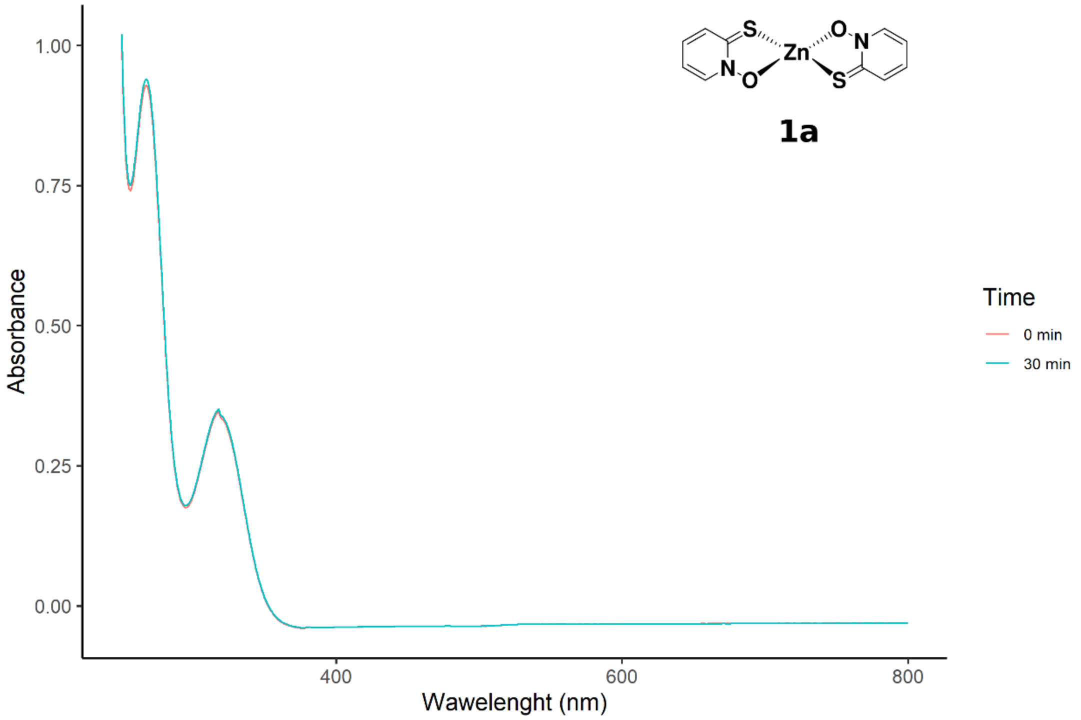

**Supplementary Figure 11: UV-vis stability spectra of zinc complex 1a in 1% DMSO/acetate buffer.**

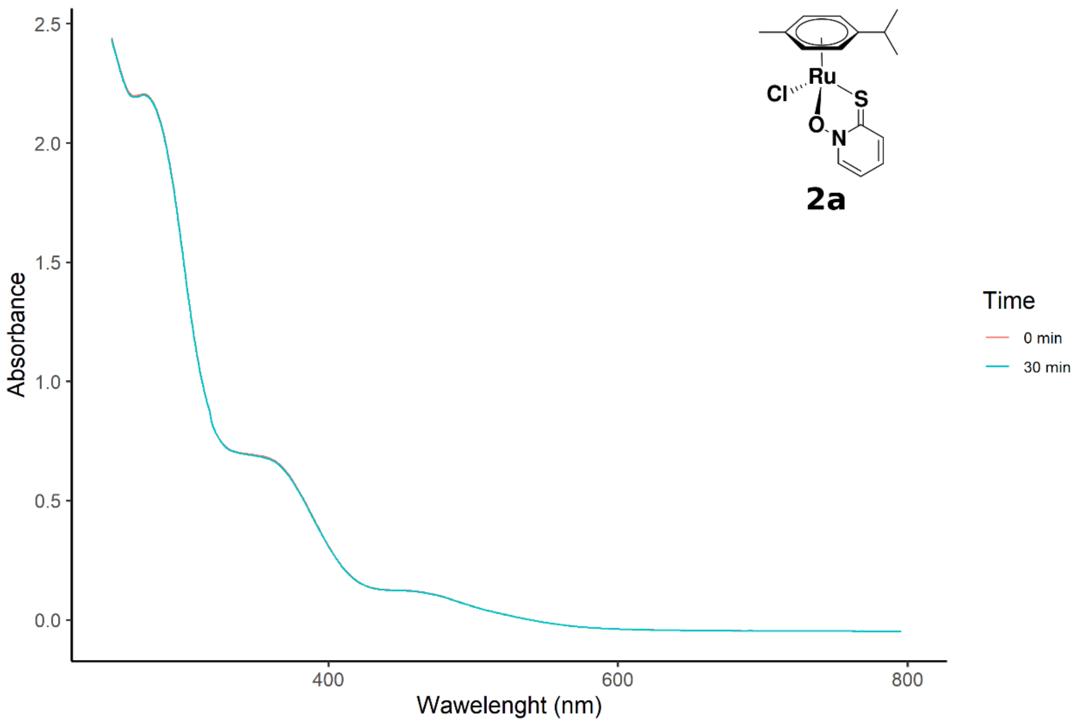

**Supplementary Figure 12: UV-vis stability spectra of ruthenium complex 2a in acetate buffer.**

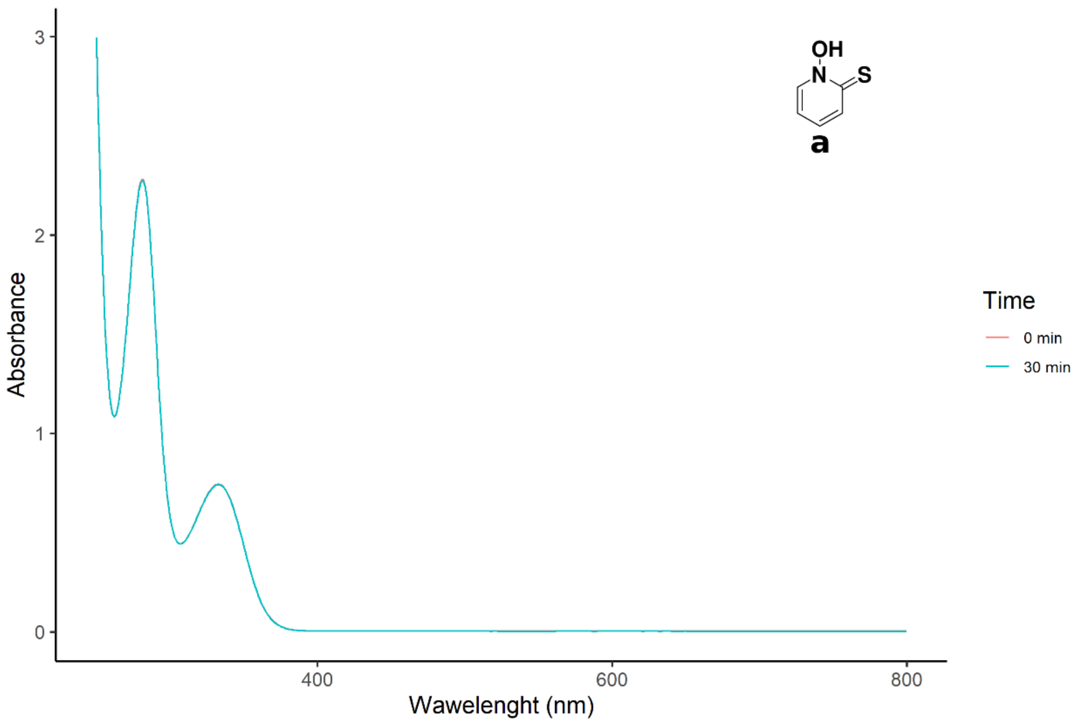

**Supplementary Figure 13: UV-vis stability spectra of ligand pyrithione a in 1% DMSO/HEPES buffer.**

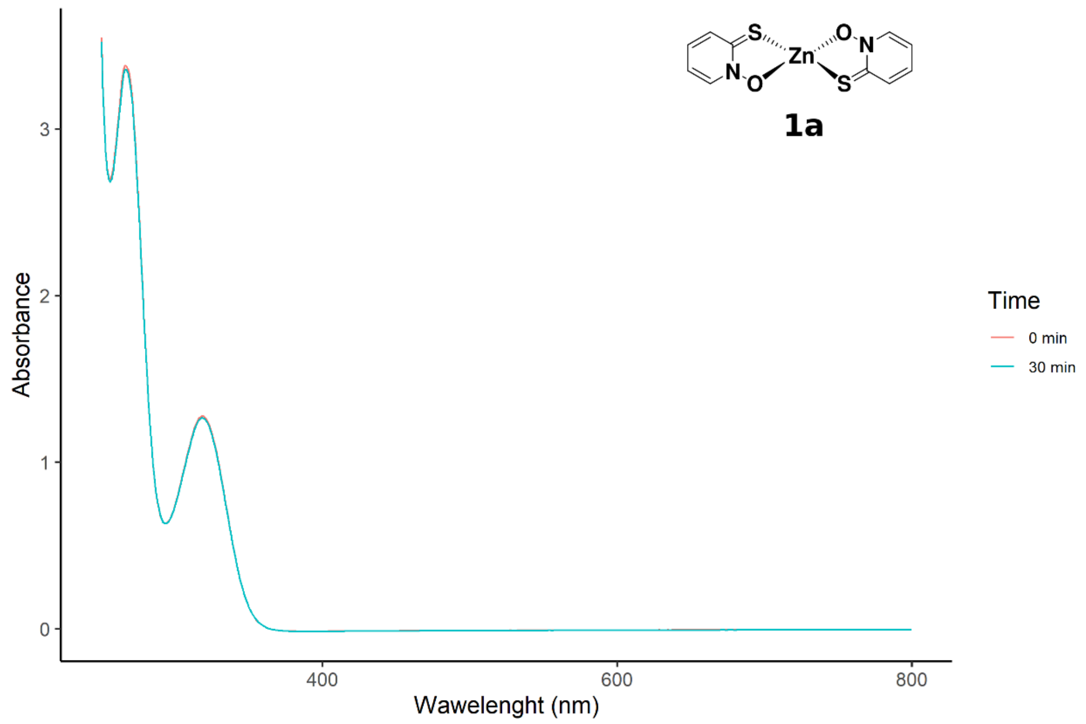

**Supplementary Figure 14: UV-vis stability spectra of zinc complex 1a in 1% DMSO/HEPES buffer.**

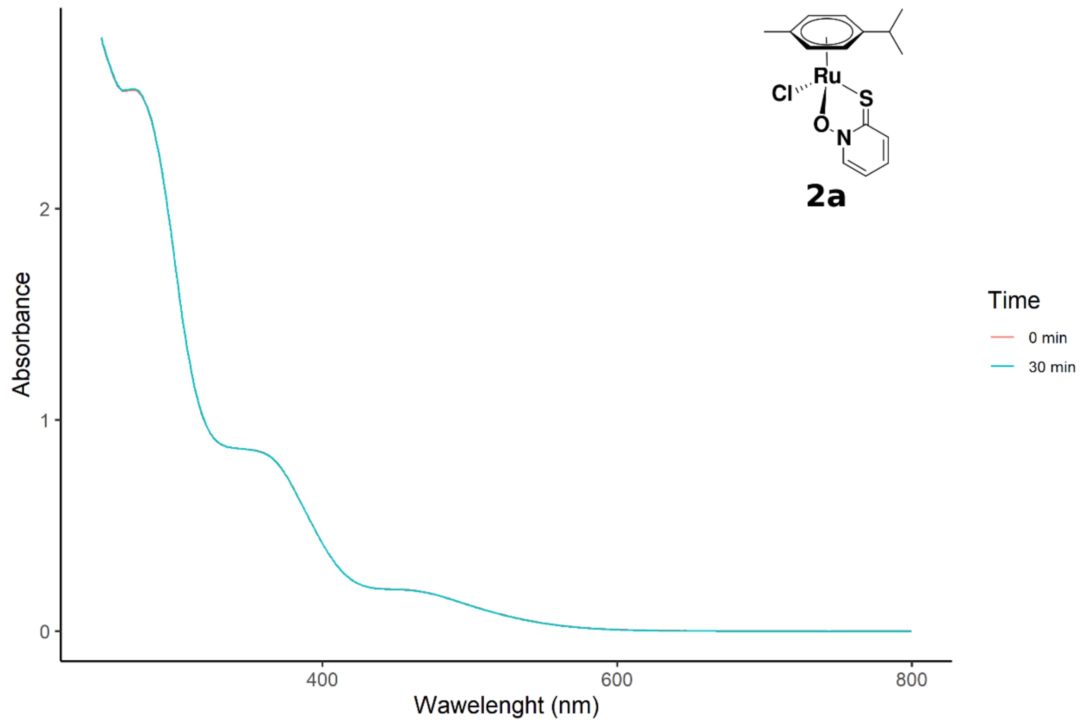

**Supplementary Figure 15: UV-vis stability spectra of ruthenium complex 2a in HEPES buffer.**

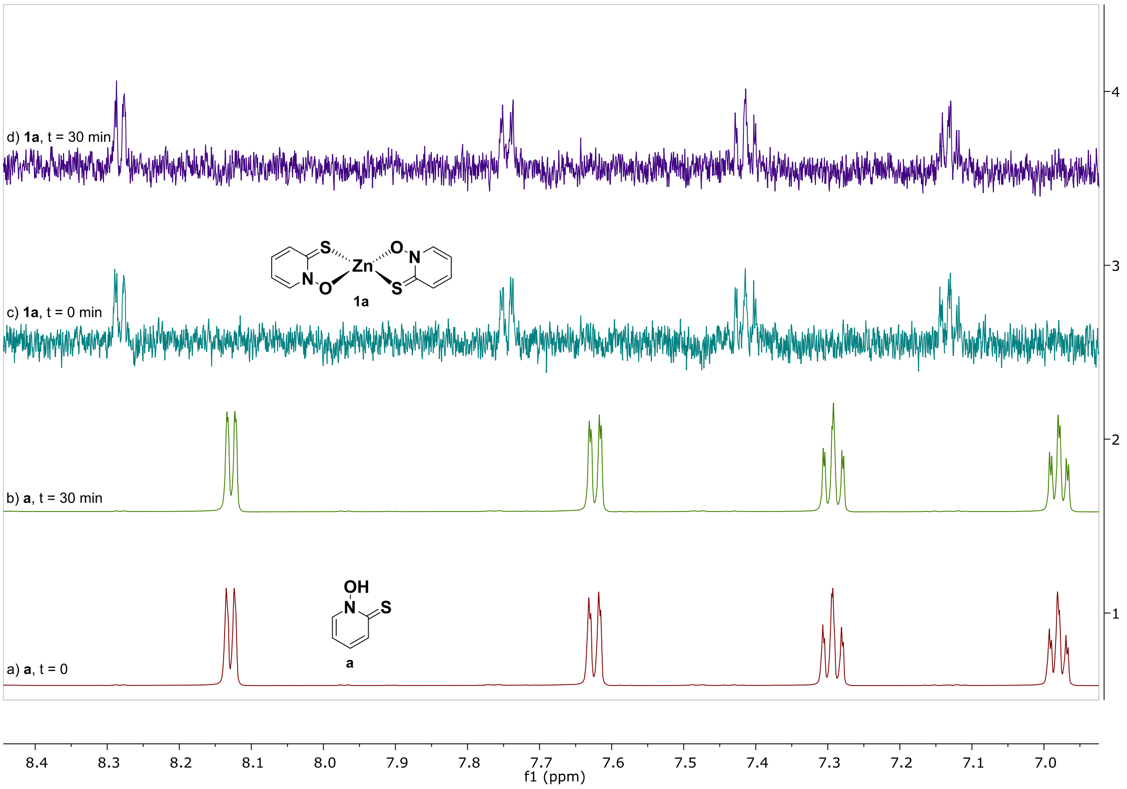

**Supplementary Figure 16: $^{1}$H NMR stability spectra of pyrithione a (a–b) and zinc complex 1a (c–d) in 1% DMSO-$d_6$/acetate buffer solution prepared in $D_2O$ recorded immediately after preparation and after 30 min.**

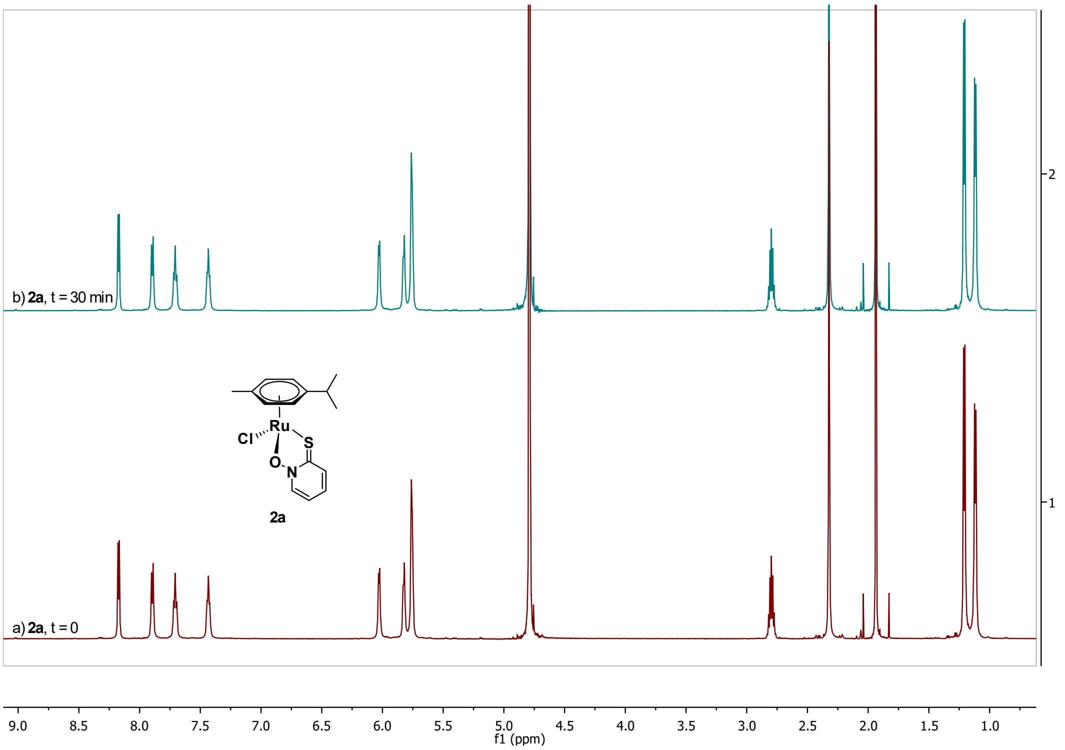

**Supplementary Figure 17: $^{1}$H NMR stability spectra of ruthenium complex with pyrithione 2a in acetate buffer solution prepared in $D_2O$ recorded immediately after preparation and after 30 min.**

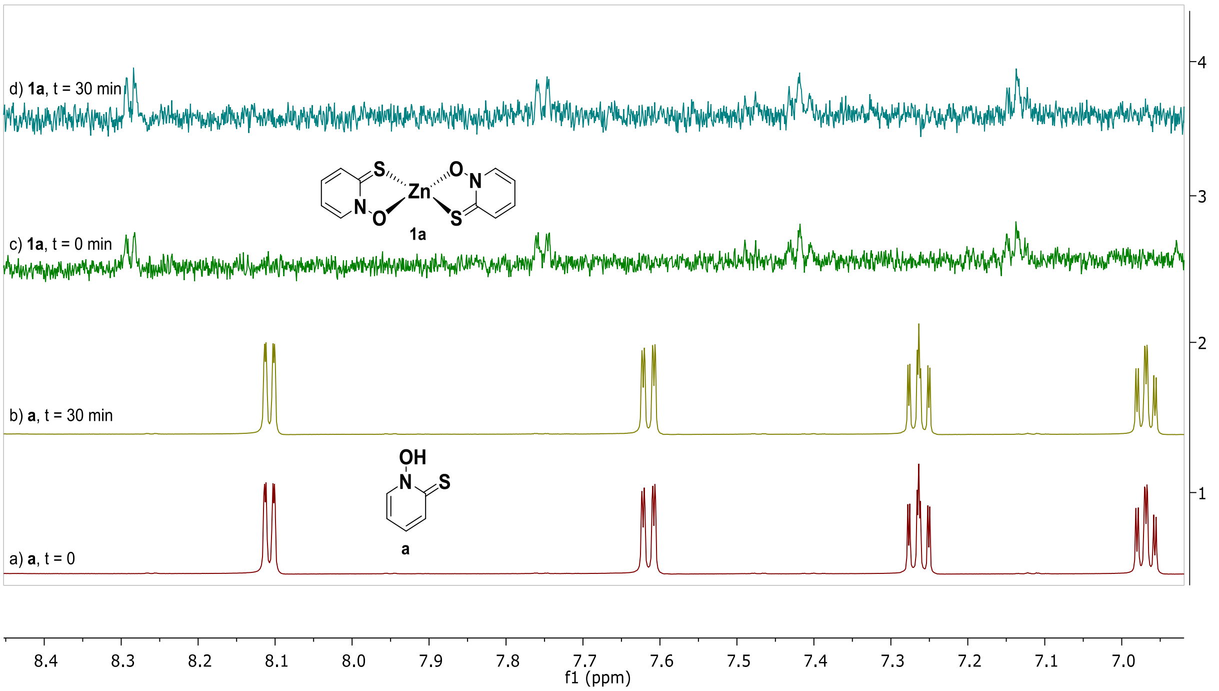

**Supplementary Figure 18: ¹H NMR stability spectra of pyrithione a (a–b) and zinc complex 1a (c–d) in 1% DMSO-$d_6$/HEPES buffer solution prepared in $D_2O$ recorded immediately after preparation and after 30 min.**

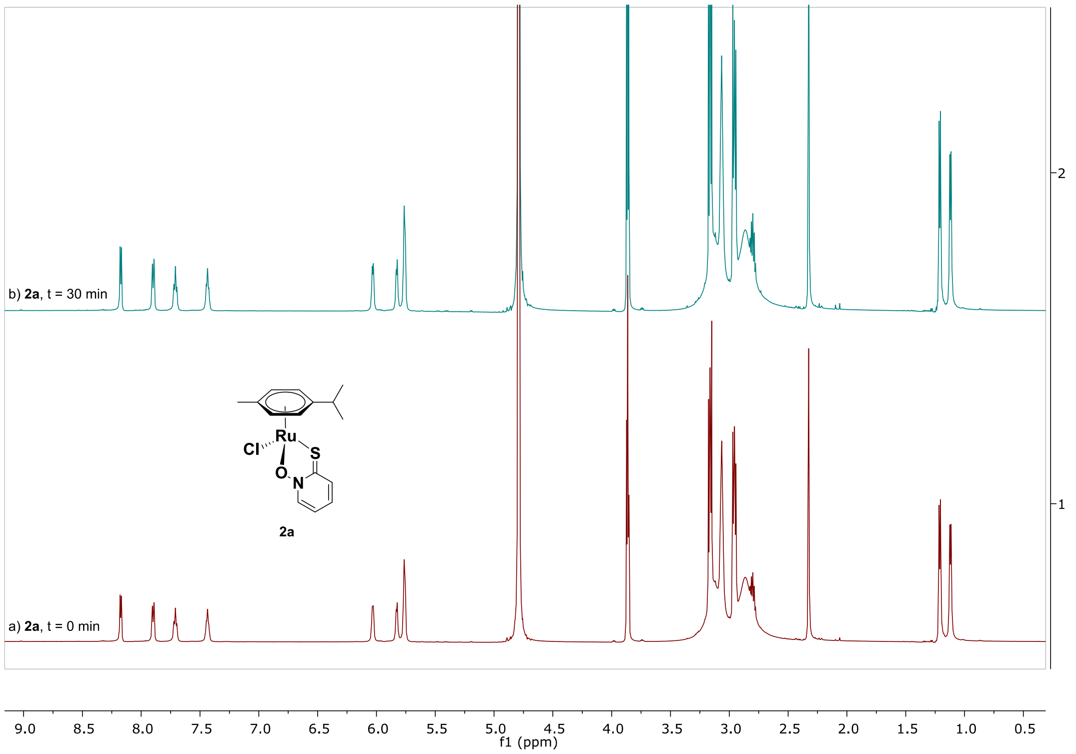

**Supplementary Figure 19: ¹H NMR stability spectra of ruthenium complex with pyrithione 2a in HEPES buffer solution prepared in $D_2O$ recorded immediately after preparation and after 30 min.**

### 6. Enzyme assays

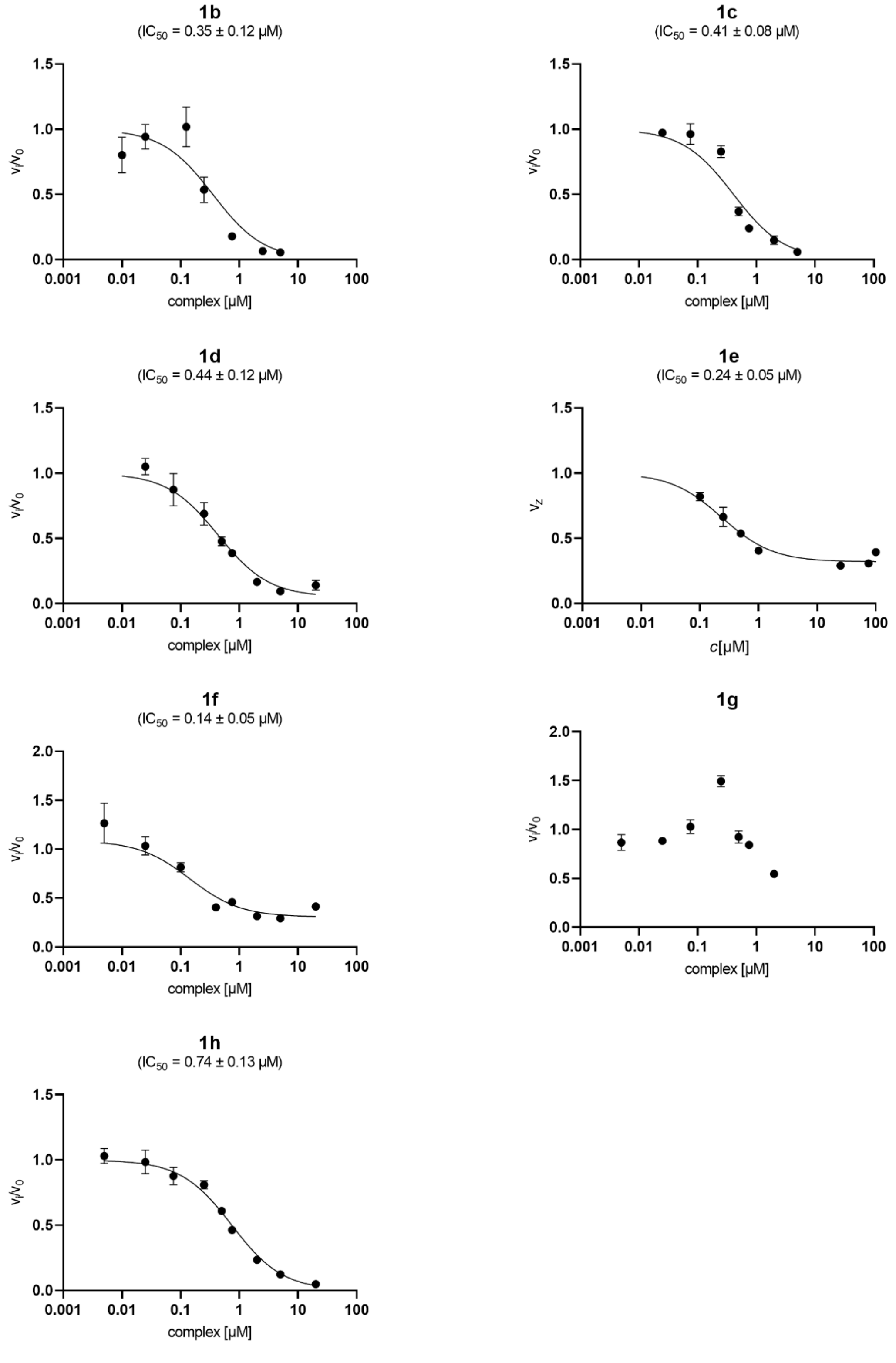

**Supplementary Figure 20**: **Results of enzyme inhibition assays for cathepsin L.** Data are mean ± s.e.m. of three measurements.

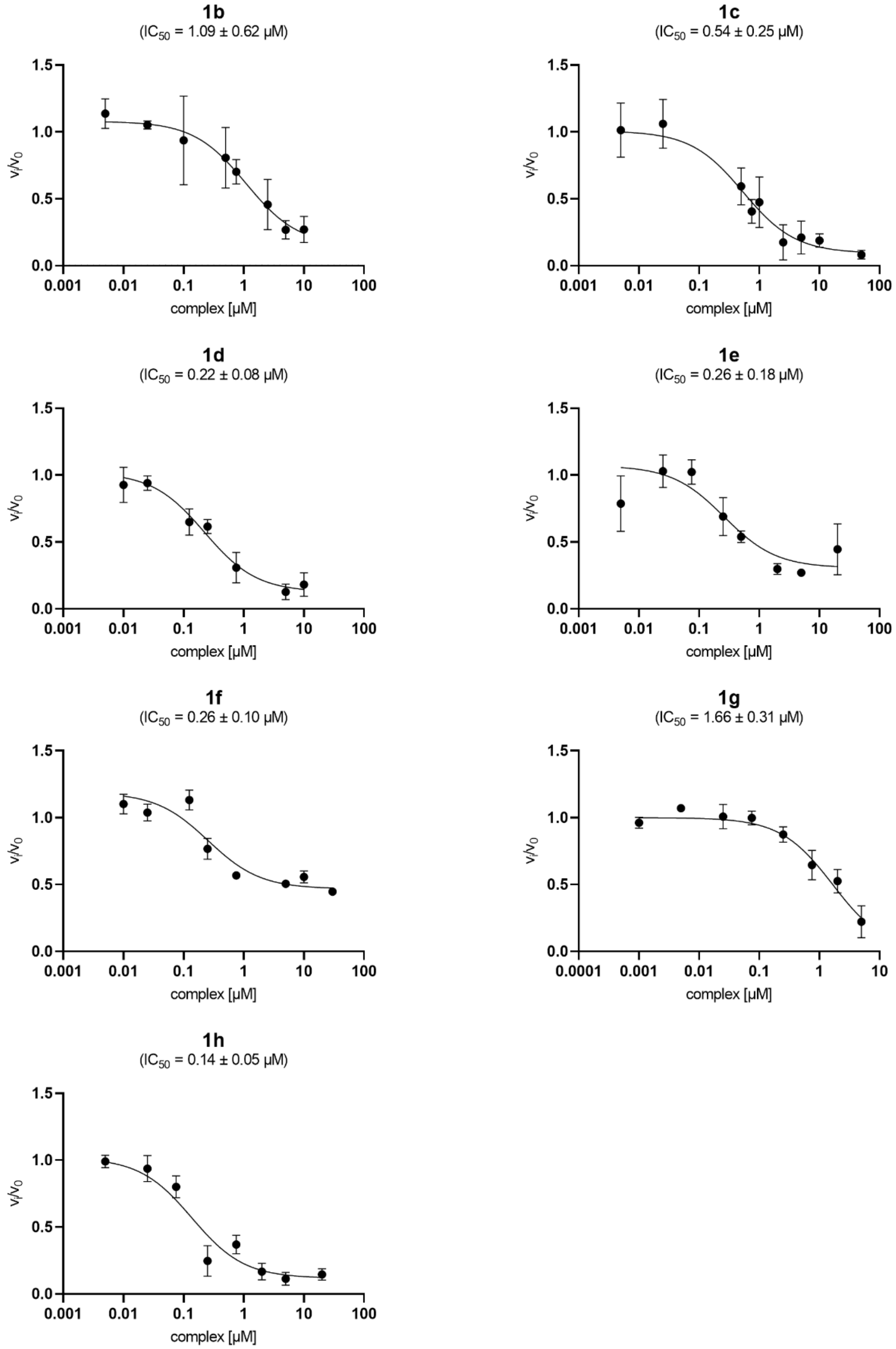

**Supplementary Figure 21**: **Results of enzyme inhibition assays for PL$^{Pro}$.** Data are mean ± s.e.m. of three measurements.

**Supplementary Table 3: Relative reaction rates for ligands tested on Cahtepsin L and $PL^{Pro}$.** Data are averages of three measurements.

| | $v_i/v_0$ at 100 µM | |
|---|---|---|
| **Ligand** | **Cathepsin L** | **$PL^{Pro}$** |
| **a (Pth)** | 1.09 | 0.88 |
| **b (3MePth)** | 1.05 | 0.50 |
| **c (4MePth)** | 1.04 | 0.60 |
| **d (5MePth)** | 1.01 | 0.77 |
| **e (6MePth)** | 1.15 | 0.82 |
| **f (1iQth)** | 0.65 | 0.66 |
| **g (2Qth)** | 0.53 | 0.28 |
| **h (3OMePth)** | 1.17 | 0.60 |

### 7. Prediction data for the key parameters in drug design

**Molecule 1**

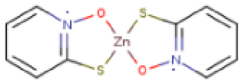

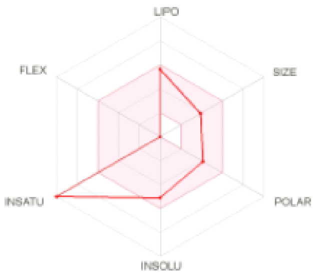

SMILES c1ccc2n(c1)O[Zn]1(S2)Sc2n(O1)cccc2

| Physicochemical Properties | |
|---|---|
| Formula | C10H8N2O2S2Zn |
| Molecular weight | 317.69 g/mol |
| Num. heavy atoms | 17 |
| Num. arom. heavy atoms | 12 |
| Fraction Csp3 | 0.00 |
| Num. rotatable bonds | 0 |
| Num. H-bond acceptors | 2 |
| Num. H-bond donors | 0 |
| Molar Refractivity | 62.55 |
| TPSA | 78.92 Å² |

| Lipophilicity | |
|---|---|
| Log $P_{o/w}$ (iLOGP) | 0.00 |
| Log $P_{o/w}$ (XLOGP3) | 4.40 |
| Log $P_{o/w}$ (WLOGP) | 2.43 |
| Log $P_{o/w}$ (MLOGP) | 1.88 |
| Log $P_{o/w}$ (SILICOS-IT) | -0.44 |
| Consensus Log $P_{o/w}$ | 1.65 |

| Water Solubility | |
|---|---|
| Log S (ESOL) | -5.10 |
| Solubility | 2.50e-03 mg/ml ; 7.87e-06 mol/l |
| Class | Moderately soluble |
| Log S (Ali) | -5.77 |
| Solubility | 5.34e-04 mg/ml ; 1.68e-06 mol/l |
| Class | Moderately soluble |
| Log S (SILICOS-IT) | -0.76 |
| Solubility | 5.47e+01 mg/ml ; 1.72e-01 mol/l |
| Class | Soluble |

| Pharmacokinetics | |
|---|---|
| GI absorption | High |
| BBB permeant | No |
| P-gp substrate | Yes |
| CYP1A2 inhibitor | Yes |
| CYP2C19 inhibitor | No |
| CYP2C9 inhibitor | Yes |
| CYP2D6 inhibitor | No |
| CYP3A4 inhibitor | No |
| Log $K_p$ (skin permeation) | -5.11 cm/s |

| Druglikeness | |
|---|---|
| Lipinski | Yes; 0 violation |
| Ghose | Yes |
| Veber | Yes |
| Egan | Yes |
| Muegge | Yes |
| Bioavailability Score | 0.55 |

| Medicinal Chemistry | |
|---|---|
| PAINS | 0 alert |
| Brenk | 1 alert: heavy_metal |
| Leadlikeness | No; 1 violation: XLOGP3>3.5 |
| Synthetic accessibility | 4.60 |

**Supplementary Figure 22**: Evaluation of physicochemical properties, pharmacokinetics, drug-likeness and medicinal chemistry friendliness of zinc pyrithione **1a**.[9]
